# Detectability of runs of homozygosity is influenced by analysis parameters and population-specific demographic history

**DOI:** 10.1101/2022.09.29.510155

**Authors:** Avril M. Harder, Kenneth B. Kirksey, Samarth Mathur, Janna R. Willoughby

## Abstract

Wild populations are increasingly threatened by human-mediated climate change and land use changes. As populations decline, the probability of inbreeding increases, along with the potential for negative effects on individual fitness. Detecting and characterizing runs of homozygosity (ROHs) is a popular strategy for assessing the extent of individual inbreeding present in a population and can also shed light on the genetic mechanisms contributing to inbreeding depression. Here, we analyze simulated and empirical data sets to demonstrate the downstream effects of program selection and long-term demographic history on ROH inference. We also apply a sensitivity analysis to evaluate the effects of various parameter values on ROH-calling results and demonstrate its utility for parameter value selection. We show that ROH inferences can be biased when sequencing depth and the distribution of ROH length is not interpreted in light of demographic history as well as program-specific tendencies. This is particularly important for the management of endangered species, as underestimating inbreeding signals in the genome can substantially undermining conservation initiatives. Based on our observations, we suggest using a combination of ROH detection tools and ROH length-specific inferences to generate robust population inferences regarding inbreeding history. We outline these recommendations for ROH estimation at multiple levels of sequencing effort typical of conservation genomics studies.

## Introduction

Climate change and expanding human land use are increasingly partitioning wild populations into smaller and smaller areas of available and suitable habitat, often leading to declining populations sizes [1,2]. Decreases in population size can lead to increased inbreeding, which has been reported to have negative fitness consequences for inbred individuals in many wild populations [3–6]. When inbreeding depression is sufficiently severe, populations can be threatened with extirpation; thus, assessing inbreeding extent is crucial for understanding and mitigating risk in small populations of conservation concern. Prior to widespread application of whole-genome sequencing strategies to non-model species, genetic estimates of inbreeding were obtained using allozyme or microsatellite data or inferred from known pedigrees [7–10]. These studies have been critically important to understanding the genetic dynamics of stable and shrinking populations and have led to increasing recognition of inbreeding depression’s prevalence and ability to affect wild population persistence [4,11]. However, applying whole genome sequencing strategies to identify runs of homozygosity (ROHs; genomic regions where both inherited haplotypes are identical) opens up lines of inquiry previously not accessible via pedigree- or microsatellite-based studies [12].

The identification and quantification of inbreeding depression, largely a result of homozygosity of strongly deleterious mutations, is critical to conservation as it guides effective management strategies for myriad species. For example, the introduction of genetically diverse individuals into declining populations to reduce the proportion of deleterious alleles present in a population is expected to reduce genetic load and, as a result, reduce extinction risks. Further, coupling genomic data with fitness outcomes has shown that longer runs of homozygosity (ROHs) are associated with higher mutational loads [13,14] and ROH analyses in historical and extant populations have identified genetic trajectories leading to species declines and extinctions [15,16]. Thus, the integration of genetic data via ROH estimation and existing demographic data can provide important clues about population stressors that support management decision making, ultimately reducing extinction risks for target species.

Estimating ROHs can provide crucial insights into populations’ evolutionary histories, but these histories can in turn affect which ROH-calling software and combination of parameter values are most appropriate. For example, the settings best suited for inferring ROHs in a small, long-isolated population experiencing high levels of inbreeding would not be suitable for individuals sampled from a large, genetically diverse population because underlying sources of error in these two scenarios are very different (*e.g.*, differences in ROH length distributions, numbers of variable sites, expected minor allele frequencies; [17]). While some studies include comparisons of results from multiple programs or parameter value combinations (*e.g.,* [18–21]), many more studies rely on default settings and do not explore the effects of varying these parameter values on their results. This has the potential to be misleading because it is always impossible to know how close the resulting estimates approximate reality, and yet ROH are an alluring metric for decision makers based on the phenomenon that they illustrate.

Here, we address the challenge of interpreting ROH population patterns using simulated and empirical genomic sequencing data to compare ROH identification patterns among different demographic histories. Here, we focus on whole-genome sequencing data, building previous studies that have examined ROH inference for data sets with lower marker densities [22–24].

Specifically, we test a wide array of setting combinations for two programs commonly used in population genomic studies—PLINK and BCFtools/RoH—and, for PLINK, apply a sensitivity analysis to evaluate the effects of parameter values on ROH inference. Based on these results, we outline a set of recommendations for ROH estimation for population genomics studies.

## Methods

### Part I: Simulated data

#### Data generation and genotype calling

We used SLiM v4.0.1 and modifications of scripts published by Stoffel et al [6] to simulate four distinct demographic scenarios: (i) a long-term large population, (ii) a long-term small population, (iii) a bottlenecked population, and (iv) a long-term declining population. Each simulation was initiated as a population of 10,000 individuals, wherein each individual consisted of a homologous pair of 30-Mb chromosomes [25,26]. Each population was simulated for 10,000 generations, followed by scenario-specific demographic changes (Fig. 1). The large population scenario involved an additional 1,000 generations at a population size of 1,000 individuals, whereas the small population scenario involved 1,000 additional generations at a population size of 250 individuals. The bottlenecked population persisted with 1,000 individuals for 900 additional generations, followed by a 50-generation interval with 50 individuals (*i.e.*, the bottleneck) and a final 50-generation interval with 250 individuals. The declining population continued for 850 generations with a population size of 1,000, followed by three 50-generation intervals of decreasing population sizes, from 500 to 250 and finally to 50 individuals.

**Fig. 1.**
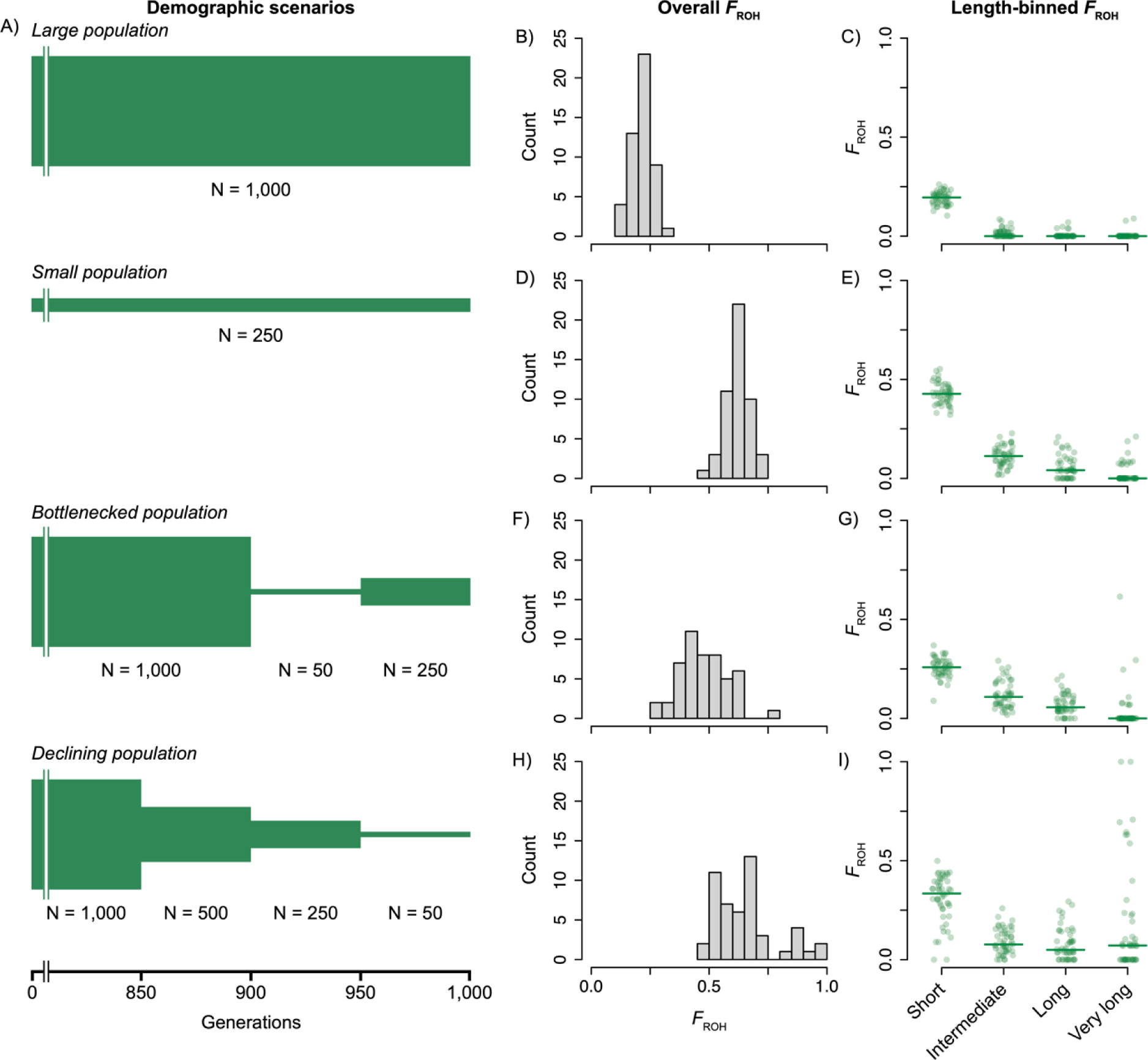
A) Diagrams describing population size changes for each of the simulated demographic scenarios. Each diagram shows the final 1,000 generations of each scenario, which was preceded in each by a burn-in of 10,000 generations with a population size of 10,000 diploid individuals. Population sizes for each interval are noted for each scenario. B,D,F,H) True overall *F*_ROH_ frequencies. C,E,G,I) Length bin-specific true *F*_ROH_ values; horizontal lines correspond to bin median values.

Recombination rate (9.15 x 10^-9^ per site per generation) and base mutation rate (1.39 x 10^-8^ per site per generation) were chosen based on estimates calculated for Tasmanian devils (*Sarcophilus harrissii*)—the focal species for our empirical tests [27,28]. Each VCF file output from SLiM was converted to FASTA sequence files using a custom script in R v4.0.3 and a haploid ancestral sequence comprising a 30-Mbp segment randomly sampled from the Tasmanian devil genome [29].

Using the known genotypes for all individuals, we generated a coordinate file for each demographic scenario recording the start and end coordinates for all true ROHs ≥ 100 kb in length (*i.e.*, ≥ 100 kb of sequential homozygous loci). We imposed this lower limit on ROH length because ROHs less than 100 kb in length likely originated in a single common ancestor approximately 500 generations ago (assuming a recombination rate of 1 cM/1 Mb; [30]), and would not be expected to influence contemporary individual fitness as strongly as more recently acquired autozygous segments [13]. This threshold is widely applied in population genetics studies of non-model species [5,31–33], and we follow this convention for all downstream analyses.

We randomly selected 50 individuals from the final generation of each population simulation and generated FASTQ read files from each of the two FASTA files representing homologous chromosomes using ART (version MountRainier-2016-06-05) [34]. We simulated 150-bp paired-end reads using the HiSeq 2500 error model to a depth of 50X per individual (*i.e.*, 25X per homologous chromosome). Each FASTQ file was quality-checked using FASTQC v0.11.9 [35]. We aligned reads to the ancestral sequence using the BWA-MEM algorithm implemented in BWA v2.0 and downsampled the resulting BAM files using SAMtools v1.17 to simulate four additional levels of coverage per individual: 5X, 10X, 15X, and 30X [36,37].

For each sorted BAM file, we called genotypes using the ‘HaplotypeCaller’ algorithm in Genomic Variant Call Format (GVCF) mode as implemented in GATK v4.1.9.0 [38]. For each level of coverage, individual GVCF files were combined using ‘CombineGVCFs’ and genotyped using ‘GenotypeGVCFs’. We applied ‘VariantFiltration’ to these VCF files in GATK to flag SNPs with low variant confidence (QualByDepth < 2), exhibiting strand bias (FisherStrand > 40), or with low mapping quality (RMSMappingQuality < 20). Finally, SNPs failing these filters and indels were removed using ‘SelectVariants.’ Although population history and genetic dataset characteristics can influence SNP filtering decisions, we chose to apply consistent filtering across all demographic scenarios to instead focus on how changing software parameters affect ROH inference. The effects of various SNP-filtering strategies on ROH inference have been addressed elsewhere [24,39].

#### ROH calling: hidden Markov model approach (BCFtools)

We applied the same ROH calling approaches to all multisample VCF files produced from the simulated data set using two of the programs most commonly applied to non-model species. First, we tested an extension of the BCFtools software package, BCFtools/RoH v1.17 [40]. This program uses a hidden Markov model to detect regions of autozygosity, requiring only a VCF file for all samples, population allele frequency information, and an optional recombination map. Because additional genetic information is not likely to be available for many wild populations, we relied on allele frequencies calculated from each of our sample sets. The main decision faced when running BCFtools/RoH is whether to estimate autozygous regions using called genotypes or genotype likelihood values. We tested the effects of this decision on ROH estimation by either including the *--GTs-only* setting to limit inference based on genotypes (hereafter, BCFtools Genotypes) or omitting it and allowing genotype likelihood values to be considered (hereafter, BCFtools Likelihoods) (Table 1). There are also two HMM options that can be set: *--hw-to-az*, which sets the transition probability of Hardy-Weinberg to autozygous state, and *--az-to-hw*, which sets the transition probability of autozygous to Hardy-Weinberg state. The default values for these two probabilities are 6.7 x 10^-8^ and 5 x 10^-9^, respectively. However, BCFtools/RoH can be run in Viterbi training mode (*--viterbi-training* option) to estimate these two probabilities for a specific data set. In total for BCFtools/RoH, we applied both methods, Genotypes and Likelihoods, using default transition probabilities and population-specific probabilities calculated using the *--viterbi-training* option and a custom R script (available in the GitHub repository for this project) to each demographic scenario and coverage level.

**Table 1.** Parameter values applied during ROH calling for both simulated and empirical data. For PLINK, a total of 486 combinations were tested. PLINK default values are underlined. ROH = run of homozygosity.

| Program | Parameter | Abbreviation | Values | Description |
| --- | --- | --- | --- | --- |
| BCFtools/RoH | --GTs-only | -- | 30 | If set, uses genotypes only and ignores likelihood values |
|  | --viterbi-training | -- | <u>1 x 10<sup>-10</sup></u> | If set, runs BCFtools/RoH in Viterbi training mode to estimate HMM transition probabilities; value sets convergence threshold |
|  | --hw-to-az | -- | <u>6.7 x 10<sup>-8</sup></u> | Transition probability from Hardy-Weinberg to autozygous state |
|  | --az-to-hw | -- | <u>5 x 10<sup>-9</sup></u> | Transition probability from autozygous to Hardy-Weinberg state |
| PLINK | --homozyg-window-het | phwh | 0, <u>1</u>, 2 | Number of heterozygous sites allowed within a window; default = 1 heterozygous site |
|  | --homozyg-window-missing | phwm | 2, <u>5</u>, 50 | Number of missing calls allowed in a window; default = 5 missing calls |
|  | --homozyg-window-snp | phws | <u>50</u>, 100, 1000 | Scanning window length in SNPs; default = 50 SNPs |
|  | --homozyg-density | phzd | <u>50</u> | Minimum density in kb (*i.e.*, maximum inverse density (kb/variant); *e.g.*, to specify minimum 1 SNP per 50 kb, set to 50); default = 50 kb |
|  | --homozyg-gap | phzg | 500, <u>1000</u> | Threshold distance in kb at which to split a ROH into two if two SNPs are too far apart; default = 1000 kb |
|  | --homozyg-window-threshold | phwt | 0.01, <u>0.05</u>, 0.1 | Proportion of overlapping windows that must be called homozygous to assign any SNP to a ROH; default = 0.05 |
|  | --homozyg-snp | phzs | 10, <u>100</u>, 1000 | Minimum number of variants that must be included in a ROH of minimum length --homozyg-kb to report it; default = 100 SNPs |
|  | --homozyg-kb | phzk | 100 | Required minimum length of sequence (in kb) spanned by number of homozygous sites specified by --homozyg-snp; default = 1000 kb |

#### ROH calling: sliding window approach (PLINK)

We tested a large number of parameter value combinations in PLINK v1.90b6.26 [41,42]. Unlike BCFtools/RoH, PLINK employs a sliding window approach to ROH identification: for each window placement, SNPs are examined for conformity to the PLINK parameter values (*e.g.*, fewer than the number of heterozygous or missing calls allowed). It is then determined, for each SNP, whether a sufficient proportion of windows overlapping that SNP are homozygous and thus, whether the SNP is determined to be located within in a ROH. PLINK has multiple parameters that can be set by the user, and we initially tested a total of 486 combinations of six of these parameters for each level of coverage (see Table 1 for list of parameters, initial values, and parameter descriptions). To explore the effects of these combinations on ROH inference, we applied an iterative approach designed by Mathur et al. [43] to explore through visualization how varying parameter settings influence overall *F*_ROH_ estimates across samples within each population and coverage level. For each iteration, we performed four steps:

1. Run PLINK with all possible combinations of different parameters to be tested, ultimately generating a matrix of parameter values (predictor variables) and inferred *F*_ROH_ (response variable) for each sample.
2. Create a linear model for each combination of parameter values (*F*_*RO* *H*_= *a* + *b*_1_*x*_1_ + ⋯+ *b*_*n*_*x*_*n*_ + *e*; where b_i_ = weight of parameter x_i_), where the values of parameter x_i_ are standardized to 1.
3. Extract standardized rank regression coefficients (SRC) from the linear regression models using the *sensitivity* package in R and visualize sensitivity indices (SRC_i_) to rank weights of each parameter [44].
4. If SRC_i_ □ 0 with little individual variation, then set the parameter *i* to the default value. If SRC_i_ is > 0 or < 0, then consider the effect described by SRC_i_ (*i.e.*, whether increasing the value of the parameter increases or decreases *F*_ROH_ and how SRC_i_ varies with called *F*_ROH_) and either define a new set of parameter values to test or select a value from the tested set.

We began the first iteration by reading the results from the initial 486 combinations of parameter values for each demographic scenario into R v4.0.3 [29]. After examining the output figures, we selected new sets of parameter values to test for each demographic scenario (S2 Table).

#### Data summarization and statistical analyses

Output files from BCFtools/RoH and the final PLINK runs were read into R for summarization and statistical analyses. We also read in true ROH data (*i.e.*, start and end coordinates for known ROHs ≥ 100 kb in length) and calculated true *F*_ROH_ values for each individual. We filtered all called ROHs to retain ROHs ≥ 100 kb in length and calculated inferred *F*_ROH_ for each demographic scenario, individual, coverage level, and method. To describe relationships between true *F*_ROH_ and called *F*_ROH_ values, we constructed a linear model for each method and coverage level with true *F*_ROH_ as the predictor variable and called *F*_ROH_ as the response variable. For each model, we calculated the 95% confidence intervals (CIs) for the slope and *y*-intercept parameters using the *confint* function in R. To determine whether true and called *F*_ROH_ values differed for each model, we tested whether the model’s *y*-intercept differed from zero and whether the slope differed from one (*i.e.*, whether the 95% CIs included zero or one, respectively). We also used the *y*-intercept and slope parameters to compare the results obtained from each BCFtools methods when using default vs. Viterbi-trained HMM transition probabilities within each demographic scenario and, for all three approaches, to determine whether each method over- or underestimated true F_ROH_ at each coverage level and how the degree of over- or underestimation changed with increasing true *F*_ROH_ values and across demographic scenarios.

To further evaluate the accuracy of each ROH identification method, we also calculated false negative (*i.e.*, failing to call a ROH present in an individual) and false positive (*i.e.*, calling a ROH that was not present in an individual) rates for called ROHs. We began by identifying overlap between true and called ROHs on a per-position basis by summing the number of bases covered by both the true ROH and called ROH(s). From this information, we calculated (i) the false negative rate: the total chromosomal length covered by true ROHs but not by called ROHs divided by the total length of true ROHs; and (ii) the false positive rate: the total chromosomal length covered by called ROHs but not by true ROHs divided by the total chromosomal length not covered by true ROHs. For each demographic scenario, method, and level of coverage, we calculated median false positive and negative rates and compared these medians and the 50% quantiles between all method and coverage level combinations to provide insight into method-specific differences in ROH calling errors.

We calculated *F*_ROH_ for ROHs in four different length bins to explore how ROH identification methods may differ in their capabilities to accurately call ROHs of different sizes. We defined length bins as:

(i) Short ROHs: ≥ 100 kb and < 500 kb in length;
(ii) Intermediate ROHs: ≥ 500 kb and < 1 Mb in length;
(iii) Long ROHs: ≥ 1 Mb and < 2 Mb in length; and
(iv) Very long ROHs: ≥ 2 Mb in length.

We examined how *F*_ROH_ for each bin changed with increasing coverage and also how patterns of over- and underestimation of *F*_ROH_ varied with increasing coverage by subtracting true *F*_ROH_ from called *F*_ROH_ for each individual. For each method, level of coverage, and length bin, we compared mean called *F*_ROH_ – true *F*_ROH_ and the 95% CI around these means (estimated using the *quantiles* function in R), with CIs < 0 indicating underestimation of true *F*_ROH_ and CIs > 0 indicating overestimation. We further explored relationships between true and called ROHs by examining how true and called ROHs overlap. We tabulated how many true ROHs each called ROH overlaps (or contains) and vice-versa for each unique combination of demographic scenario, ROH detection method, coverage level, and ROH length bin.

### Part II: Empirical data

#### Data curation and genotype calling

To test the effects of program and parameter value selection on identifying ROHs from empirical data, we analyzed publicly available whole genome sequencing data for a species of conservation concern, the Tasmanian devil (BioProject PRJNA549794 in NCBI’s Sequence Read Archive; [45]). From the full data set, we selected the 15 individuals from this data set with the highest number of reads. The accession numbers and relevant metadata for each set of sequences are provided in S2 Table. Adapters and low-quality bases were trimmed from raw sequences using Trim Galore v0.6.6 [46], and cleaned reads were mapped to the chromosome-level mSarHar1.11 *S. harrissii* reference genome (NCBI GenBank accession GCA_902635505.1) using BWA-MEM [36].

We used Qualimap v2.2.1 to determine mean coverage per individual from each sorted BAM file [47]. These results were used to calculate the downsampling proportions required to approximate 5X, 10X, 15X, and 30X coverage for each individual. Following downsampling, BAM files were processed in the same manner as for the simulated data, with additional SNP filtering criteria applied in VCFtools v0.1.17 [48], including filtering SNPs within 5 bp of indels and requiring minor allele frequencies ≥ 0.05 and < 20% missing data across individuals.

#### ROH calling and sensitivity analyses

We called ROHs from the final multisample VCF files using the same approaches as for the simulated data. We called ROHs in two ways, (i) using BCFtools/RoH (*i.e.*, relying on genotypes or on genotype likelihood values and using default or Viterbi-trained HMM transition probabilities) and (ii) testing 486 parameter combinations in PLINK at each level of coverage and visualizing the effect of altering each parameter following the same sensitivity analysis process described above. Parameter values for all iterations tested are provided in S1 Table with additional details provided for the empirical data in the Supporting Information.

#### Data summarization and statistical analysis

Output files from BCFtools/RoH and the final PLINK runs for the empirical data were read into R for summarization and statistical analyses. Following the approach we used for the simulated data, we filtered all called ROHs to retain ROHs ≥ 100 kb in length and calculated inferred *F*_ROH_ for each individual in each scenario, coverage level, and method. We also calculated *F*_ROH_ for ROHs in four different length bins, where length bins were defined as described above for the simulated data. To compare results across methods, coverage levels, and ROH lengths, we calculated mean length-specific *F*_ROH_ values and compared the 95% CIs around these means among methods and coverage levels.

## Results

### Part I: Simulated data

#### Data collection and curation

For each simulated population, all analyses were based on 50 randomly sampled individuals. For the declining population, extremely low heterozygosity in two individuals precluded estimation of HMM transition probabilities via Viterbi training, and all downstream analyses for this demographic scenario were based on the remaining 48 individuals. Mean heterozygosity (□ SD) varied across scenarios: large population mean = 3.17 x10^-5^ □ 1.26 x 10^-5^; bottlenecked population mean = 3.05 x 10^-5^ □ 5.81 x 10^-6^; small population mean = 2.87 x 10^-5^ □ 1.22 x 10^-5^; declining population mean = 2.44 x 10^-5^□ 8.81 x 10^-6^. Mean *F*_ROH_ generally increased with decreasing mean heterozygosity: large population mean = 0.21; bottlenecked population mean = 0.48; small population mean = 0.62; declining population mean = 0.65 (Fig. 1B,D,F,H). For all demographic scenarios, length-specific *F*_ROH_ mostly decreased with increasing ROH length with some notable exceptions in the bottlenecked and declining population scenarios (Fig. 1C,E,G,I). For these two final sets of samples, a few individuals had high proportions of their genomes located in very long ROHs (Fig. 1G,I), consistent with very recent inbreeding events in each population. Following downsampling and SNP filtering, the final mean coverage across all demographic scenarios was 4.87, 9.77, 14.69, and 29.20, and 44.99 for the 5X, 10X, 15X, 30X, and 50X VCF files, respectively.

#### ROH calling

Before comparing results from the three main methods—BCFtools Genotypes, BCFtools Likelihoods, and PLINK—we examined the effect of using either default or Viterbi-trained HMM transition probabilities on the results produced by each BCFtools method. Within each method, we compared overall *F*_ROH_ values obtained when using default HMM transition probabilities (default values: *--hw-to-az* = 6.7 x 10^-8^, *--az-to-hw* = 5 x 10^-9^) against those calculated when using probabilities estimated via Viterbi training (mean values across all groups: *--hw-to-az* = 9.2 x 10^-7^, *--az-to-hw* = 3.8 x 10^-6^; S3 Table). (In practice, mean probabilities were calculated and applied for each demographic scenario and BCFtools method combination, but note the small magnitude of variation in Viterbi-trained probabilities across all groups relative to the difference between these probabilities and the program defaults.) The slope and intercept parameters never significantly differed between the two sets of results across all demographic scenarios and coverage levels, but for most comparisons, the slope and intercept parameters of the model built using Viterbi-trained results were closer to 1 and 0, respectively, than for the default values model (S4 Table; S1-4 Figs.). Using Viterbi-trained transition probabilities slightly increased false positive rates but also decreased false negative rates (S5-8 Figs.). Because of these slight, if not significant, improvements in overall *F*_ROH_ estimation accuracy and false negative rates, we chose to proceed with Viterbi-trained HMM transition probabilities for both BCFtools methods and present only these results below (we also describe one additional benefit of Viterbi-trained probabilities below).

We used our simulated data set and linear models to determine whether each approach tends to over- or underestimate true *F*_ROH_. All three methods underestimated *F*_ROH_, with all model intercepts across coverage levels negative and different from zero, except for PLINK when calling ROHs in the large population at 5X coverage (*i.e.*, no other 95% CIs for intercepts included zero; Fig. 2; S5 Table). Within each demographic scenario, the three methods produced similar results with respect to overall *F*_ROH_ accuracy. For example, within all four scenarios, intercepts did not differ among methods or coverage levels, with two exceptions at 5X coverage when using PLINK: intercepts differed between this group and others within the large, declining, and majority of coverage levels for small populations. Perhaps the most visualizing striking points of comparison across demographic scenarios are the slope estimates specific to each.

**Fig. 2.**
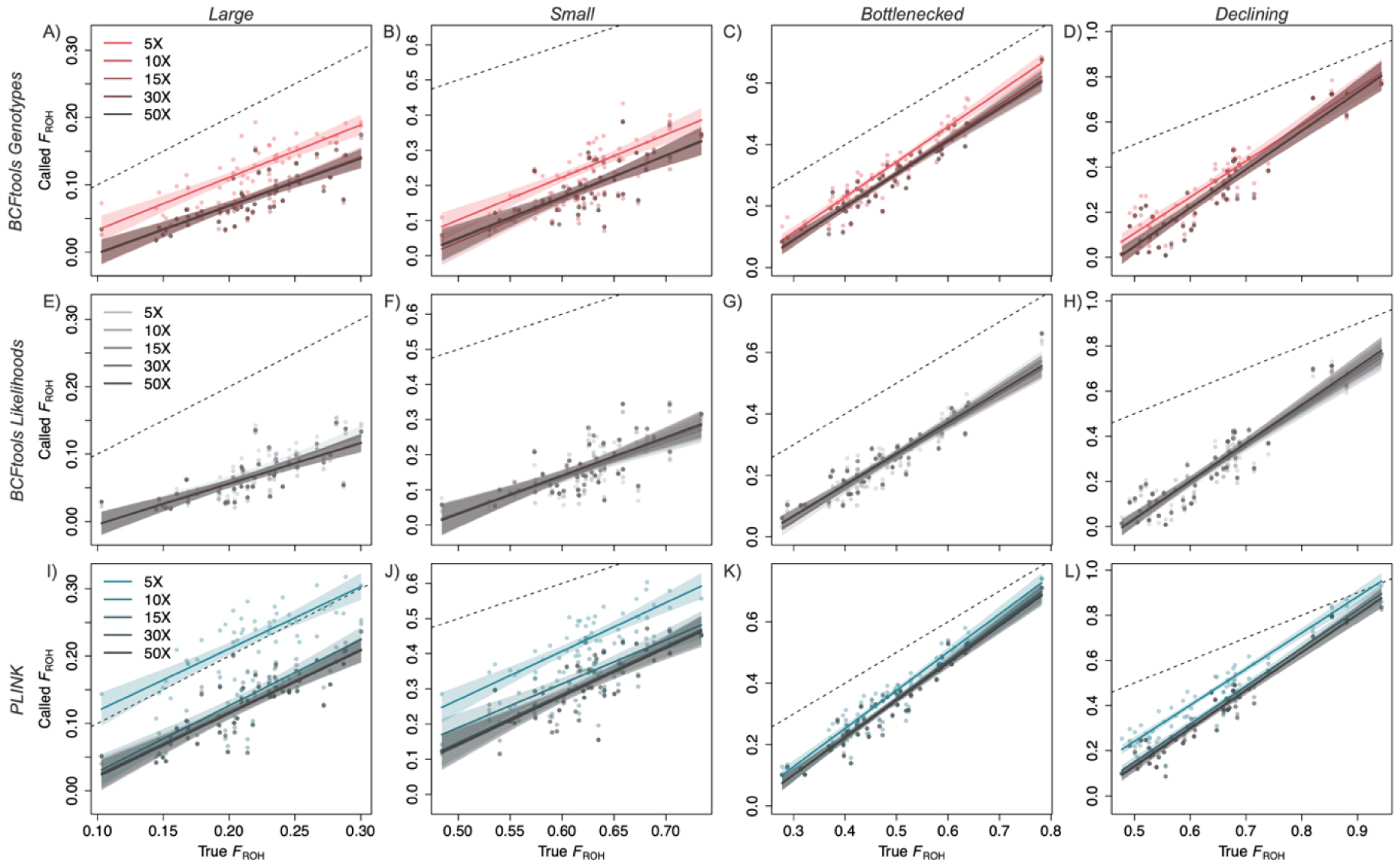
The relationship between true and inferred *F*_ROH_ values depends on inference method and population demographic scenario. Each regression line represents linear model results for a single level of coverage with the shaded areas representing 95% confidence intervals. Each point represents data for a single simulated individual. Dashed line is 1:1 line and *x*- and *y*-axes are consistent within each demographic scenario. Note the differing slopes across demographic scenarios (*e.g.*, among panels A-D) and differing overall accuracies across methods (*e.g.*, differing distances between regression lines and 1:1 line among panels D, H, and L).

Within each method, demographic scenarios were consistently ordered by slope estimates, with the declining population having the largest slope estimate (for BCFtools, all slopes > 1 with no 95% CIs including one), followed by the small, bottlenecked, and large populations (for the large population, all BCFtools slopes < 1, all PLINK slope 95% CIs include one; Fig. 2; S5 Table).

Thus, accuracy of overall *F*_ROH_ estimates calculated for the declining population varied the most with respect to true *F*_ROH_ (*i.e.*, of all demographic scenarios, slope parameter estimates for this population most greatly differed from one), followed by the large population. With respect to true *F*_ROH_, accuracy went up with increasing true *F*_ROH_ for the declining population and went down with increasing true *F*_ROH_ for the large population.

We calculated false negative (*i.e.*, failing to call a ROH present in an individual) and false positive (*i.e.*, calling a ROH that was not present in an individual) rates to further assess each method’s accuracy. With respect to false positive rates, PLINK performed poorly relative to the other two methods, with false positive rates consistently higher than BCFtools Genotypes and BCFtools Likelihoods (Fig. 3A). Across all demographic scenarios and coverage levels, median false positive rates ranged from 6.14 x 10^-7^ to 0.031 for BCFtools Genotypes, from 4.54 x 10^-8^ to 0.007 for BCFtools Likelihoods, and from 0.008 to 0.112 for PLINK. For all three methods, increasing coverage to 10X corresponded to decreasing false positive rates, but rates did not continue to improve at coverages above 10X (*i.e.*, 50% quantiles around median values overlap at 10-50X within each method and demographic scenario; results for all coverage levels provided in S9-10 Figs.). Variation in false positive rates among samples at each coverage level was smallest for BCFtools Likelihoods, followed by BCFtools Genotypes, with PLINK showing the greatest variation across samples (summary statistics provided in S6 Table).

**Fig. 3.**
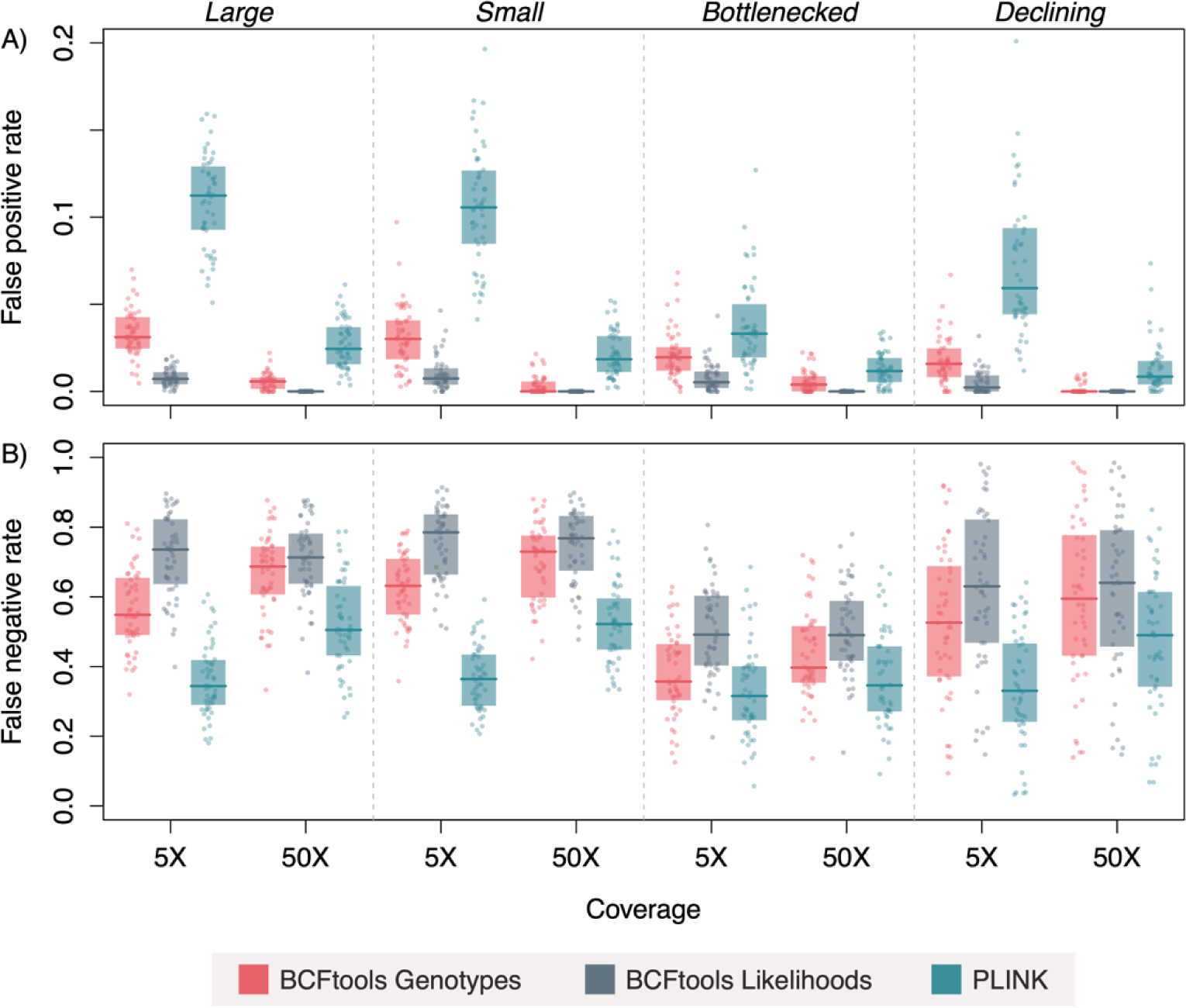
PLINK outperforms BCFtools with respect to false negative rates, but underperforms with respect to false positive rates. A) False positive (*i.e.*, incorrectly calling a base position as being located in a ROH) and B) false negative (*i.e.*, failing to identify a base position as being located in a ROH) rates across demographic scenarios and methods. Horizontal lines indicate median values and shaded boxes are 50% quantiles. Note difference in scale of *y*-axis between panels A and B. Both BCFtools approaches outperform PLINK with respect to false positive rates but the reverse is true for false negative rates. Increasing coverage corresponds to decreasing false positive rates and to increasing false negative rates. Values displayed for 5X and 50X coverages; data for all coverage levels presented in Fig. S9-10.

Patterns in false negative rates were generally in the opposite direction and magnitude to those we observed with false positives: both BCFtools methods performed poorly relative to PLINK, with BCFtools Genotypes producing slightly lower rates (median range across all demographic scenarios and coverage levels = 0.36-0.74) than BCFtools Likelihoods (median range = 0.49-0.78; Fig. 3B). PLINK exhibited lower false negative rates than the BCFtools approaches (median range = 0.32-0.53) and less variation among samples at each coverage level.

All three methods produced false negative rates that increased slightly at 10X coverage relative to 5X, with rates leveling off at 10X and higher coverage levels. Examples of false negative and false positive scenarios can be seen in S11 Fig., which illustrates a full chromosome of true and called ROHs for one exemplar individual from each demographic scenario.

We also examined how true and called values of *F*_ROH_ varied for ROHs of different lengths. For the simulated data, all three methods almost always underestimated the proportion of the genome located in short ROHs across demographic scenarios, with the 95% CI less than zero for all tests other than PLINK at 5X coverage (Fig. 4; all levels of coverage presented in S12-15 Figs.). For longer ROHs, results were variable and dependent on demographic scenario. For example, all three methods appeared to approach accuracy for longer ROHs in the large population, but this is likely due to the relative paucity of longer ROHs in this demographic scenario (Fig. 1). For the bottlenecked and declining populations (wherein some individuals had very long ROHs), both PLINK and BCFtools Genotypes tended to overestimate *F*_ROH_ for very long ROHs. PLINK generally produced the highest overestimates of *F*_ROH_ and the most variation across samples of the three approaches, followed by BCFtools Genotypes. Finally, one coverage-related trend emerged across ROH length categories, methods, and demographic scenarios, with *F*_ROH_ estimates calculated at 5X coverage often exceeding estimates calculated at higher coverage levels. Across all length bins and demographic scenarios, individual estimates of length-specific *F*_ROH_ calculated at 5X were greater than those calculated at 10X for 37%, 53%, and 53% of BCFtools Likelihoods, BCFtools Genotypes, and PLINK estimates, respectively.

**Fig. 4.**
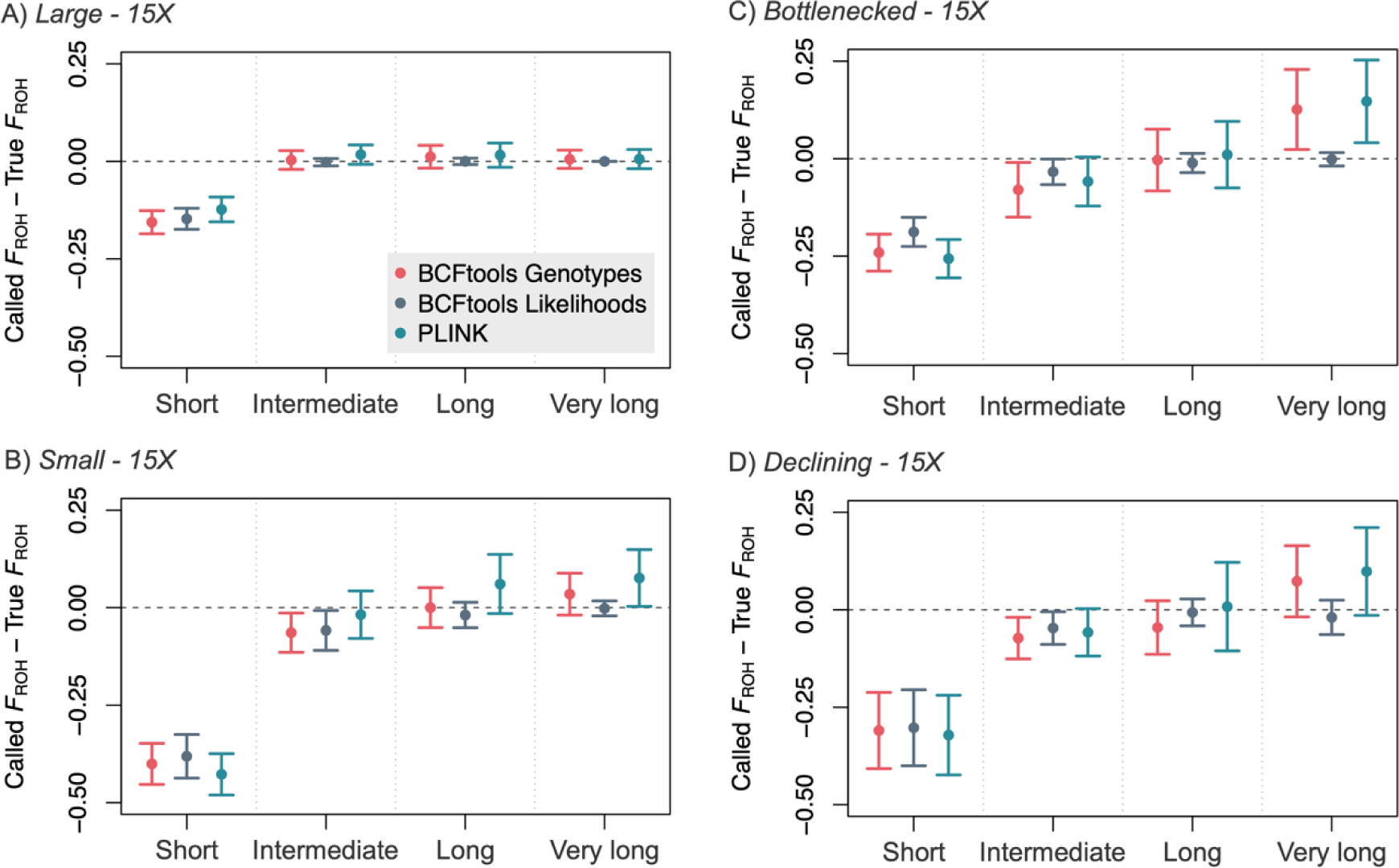
Increasing true ROH length corresponds to increasing detection. Called *F*_ROH_ – True *F*_ROH_ by length bin and demographic scenario at 15X (results for all coverage levels presented in Fig. S12-15). For BCFtools Genotypes and PLINK, *F*_ROH_ for short ROHs is consistently underestimated whereas *F*_ROH_ for very long ROHs is overestimated when these ROHs are present. BCFtools Likelihoods does not overestimate ROHs in any length bin.

To further investigate how called ROHs correspond to true ROHs, we identified regions of overlap between true and called ROHs within each individual and at each coverage level using a unique identifier for each true and called ROH. We found no instances of true ROHs being split into multiple called ROHs, but multiple true ROHs were often lumped together into a single called ROH. This pattern held true for all three methods at most coverage levels (Fig. 5; S16 Fig.). For BCFtools Genotypes and PLINK, increasing coverage did not appear to ameliorate this problem (*i.e.*, the mean number of true ROHs lumped into a single called ROH changed very little with increasing coverage; Fig. 5C,D). However, for BCFtools Likelihoods, the number of true ROHs contained in a single called ROH decreased with increasing coverage, reaching a 1:1 ratio at 30X. Across methods, the mean number of true ROHs combined into a single called ROH increased with increasing ROH length, except for BCFtools Likelihoods at coverage levels ≥ 30X (S16 Fig.). This lumping can be seen in Fig. 5B.

**Fig. 5.**
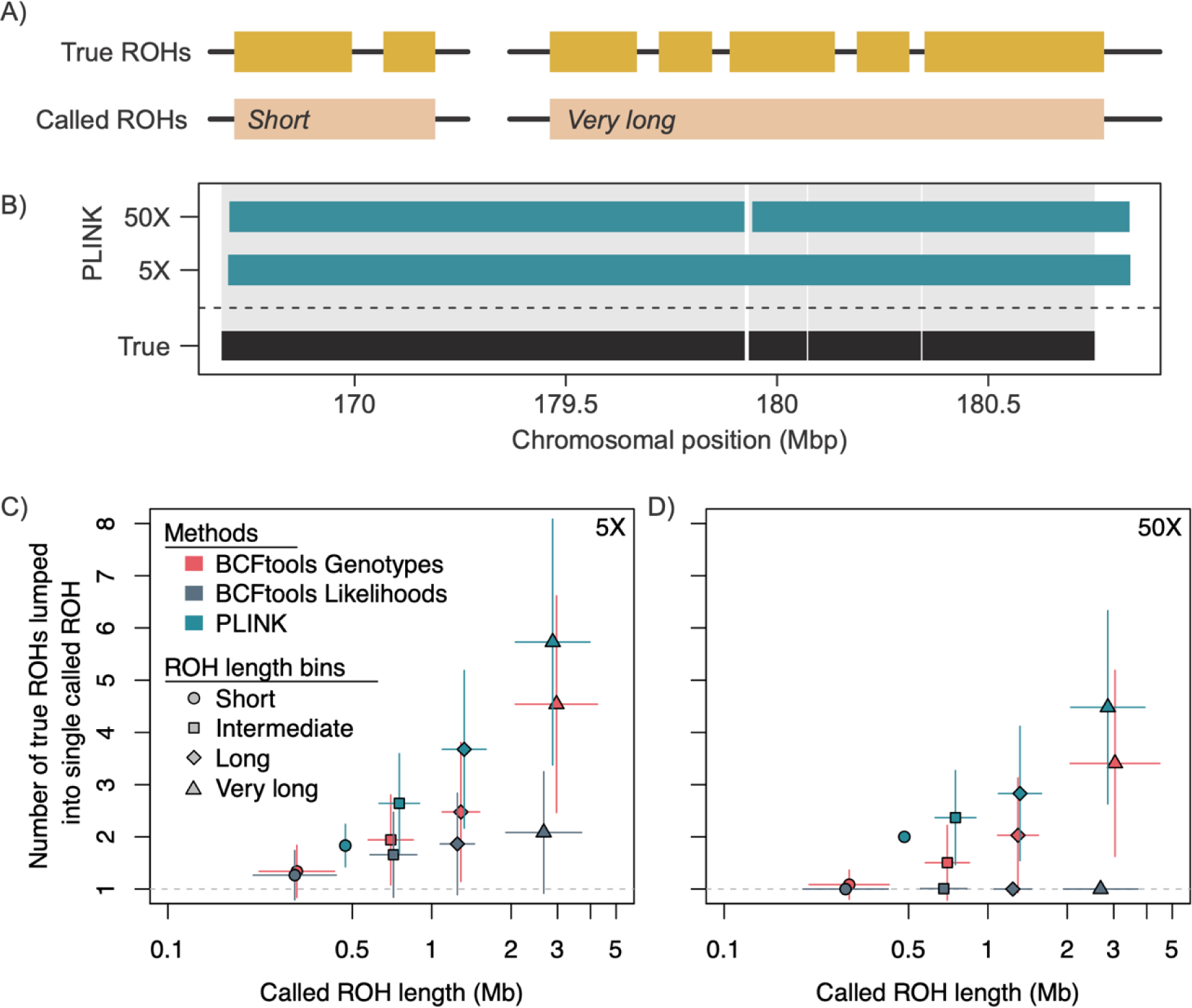
All three methods tested combine multiple true ROHs into single called ROHs, with increasing coverage only providing improvements for BCFtools Likelihoods. A) Diagram illustrating this lumping issue. B) Examples of this issue at 5X and 50X in a single simulated individual drawn from the small population demographic scenario. C) Number of true ROHs combined into a single called ROH for ROHs of varying lengths when called by all three methods at 5X and (D) at 50X in the small population (results for all coverage levels and demographic scenarios provided in Fig. S16). Points correspond to mean values and vertical and horizontal error lines indicate 95% confidence intervals. Dashed horizontal line corresponds to *y* = 1 (a 1:1 relationship between numbers of true and called ROHs).

### Part II: Empirical data

#### Genotype and ROH calling results

For the 15 sets of reads we downloaded from NCBI, the mean number of reads per sample was

9.75 x 10^8^. Read mapping rates to the mSarHar1.11 *S. harrissii* reference genome were high, with an average of 95.4% of reads mapped and properly paired. For the final sets of filtered SNPs (n = 1,532,598), average depth across samples was 48.43 for the full coverage set (*i.e.*, not downsampled) and 6.37, 11.84, 16.63, 30.75 for the 5X, 10X, 15X, and 30X downsampled sets, respectively (S2 Table). Given the modest improvements in *F*_ROH_ accuracy conferred by using Viterbi-trained HMM transition probabilities rather than the default values for the simulated data, we calculated Viterbi-trained values for both BCFtools methods applied to the empirical data.

Across methods, *F*_ROH_ estimated at 5X coverage was significantly higher than *F*_ROH_ estimates at all higher levels of coverage (95% CIs did not overlap, Fig. 6A-C). At 5X coverage, *F*_ROH_ estimates differed significantly among all three methods, with the lowest estimate produced by PLINK and the highest by BCFtools Genotypes. For all higher levels of coverage, *F*_ROH_ estimates produced by the two BCFtools methods did not differ from one another but estimates from both methods differed from those produced by PLINK, with PLINK again providing lower estimates.

**Fig. 6.**
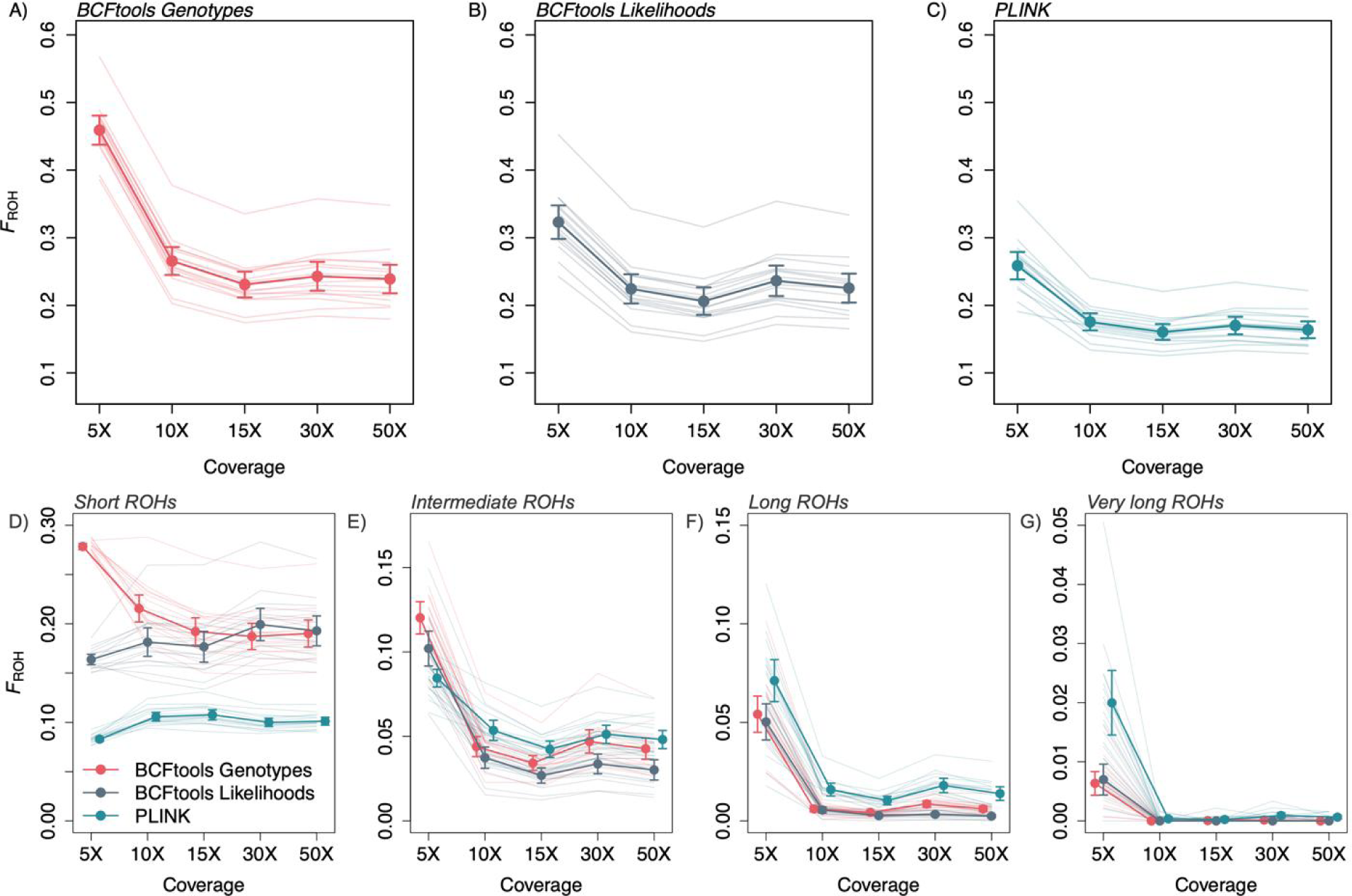
When applied to the empirical data set, the three ROH calling methods differ greatly in their inferences, particularly at 5X coverage. A-C) Overall *F*_ROH_ and (D-G) length bin *F*_ROH_ results for each method and level of coverage, with means and 95% confidence intervals indicated by points and vertical lines, respectively. Lighter background lines indicate results for individual samples.

When comparing how the three methods estimated length-specific *F*_ROH_ values, patterns varied across ROH length categories. For short ROHs, PLINK produced much lower *F*_ROH_ estimates then the two BCFtools methods across all coverage levels, with differences among the three methods significant (*i.e.*, non-overlapping 95% CIs) at 5X coverage (Fig. 6D). For longer ROHs, results of the three methods were generally concordant, with all methods producing higher estimates of *F*_ROH_ at 5X than at higher coverages (Fig. 6D-G). However, PLINK produced much higher estimates of *F*_ROH_ for very long ROHs at 5X than the other two methods, with a mean value more than double those inferred by the BCFtools methods (Fig. 6G).

## Discussion

In this study, we systematically analyzed the discrepancies that arise in ROH detection when populations with varied demographic histories are analyzed and highlight effects on downstream interpretations associated with these differences. This is important because the demographic scenarios affected the accuracy, with declining populations showing the highest variance in *F*_ROH_ estimation relative to true *F*_ROH_ values. Additionally, ROH detection accuracy varied with sequencing coverage; improvements were observed up to a 10X coverage threshold beyond which the benefits plateaued. Our findings emphasize the impact of choosing appropriate computational parameters and underscore the challenges in standardizing ROH detection methods values (*e.g.*, Saremi et al., 2019; Grossen et al., 2020; von Seth et al., 2021; Mueller et al., 2022), highlighting the necessity for robust, systematic comparisons to enhance reproducibility and reliability in genomic studies.

We demonstrated the exploratory utility of the sensitivity analysis process we followed to select parameter values for our data (see the Supporting Information for corresponding information for the empirical data). This process is important because disparate sequencing data characteristics are likely to require different parameter values, meaning that it may not be appropriate to use the values we used herein when analyzing other data. For example, studies that use fewer SNPs (*e.g.*, populations that are less genetically diverse, studies with reduced sequencing efforts) should test the effects of altering the minimum SNP density required on ROH inference results. Sensitivity analysis provides a relatively quick and convenient way to visualize how different parameter values affect *F*_ROH_ estimates for an entire data set and the degree of variation in those effects across individuals.

### Inference methodology biases F_ROH_ values

Between the two BCFtools methods when considering identified ROHs of all lengths, Genotypes produced more accurate overall *F*_ROH_ estimates than Likelihoods, with *F*_ROH_ estimates from Likelihoods also increasingly diverging from the true *F*_ROH_ value with increasing true *F*_ROH_ (Fig. 1A,B). For populations expected to have considerable variation in *F*_ROH_ among individuals (*e.g.*, a population that has remained somewhat small for an extended period of time with evidence of recent immigration), applying the BCFtools Likelihoods approach could result in increasingly skewed values for the individuals with the highest levels of inbreeding. For example, using the linear model parameters estimated for 15X coverage, an individual with a true *F*_ROH_ of 0.10 would be assigned an inferred *F*_ROH_ of 0.01 (difference = −0.09), whereas an individual with a true *F*_ROH_ value of 0.40 would be assigned 0.23 (difference = −0.17). This could be particularly problematic when dealing with species or populations of conservation concern because the individuals with the highest true *F*_ROH_ also have the largest magnitude of error, meaning that concerning signals of inbreeding could go undetected.

In contrast to the underestimations produced by the BCFtools/RoH methods, the sliding window approach implemented in PLINK overestimated *F*_ROH_. This was particularly evident at 5X coverage where *F*_ROH_ estimates differ more from their true values than any other method and coverage level combination in our study (Fig. 1C). However, at coverages above 5X, PLINK produced better estimates than either BCFtools approach (*i.e.*, in our linear models, intercepts for PLINK at 10X-50X are closer to zero than for either BCFtools method and 95% CIs for these parameter estimates do not overlap with any BCFtools intercept 95% CIs). In the context of endangered species conservation, small overestimations of *F*_ROH_ may be more desirable than underestimations because these are likely to be more conservative (*i.e.*, indicating more close inbreeding than is present in reality) in many situations. Importantly though, as with BCFtools Likelihoods, *F*_ROH_ estimates diverged from true *F*_ROH_ at increasing values of true *F*_ROH_. However, these values diverged at a much lower rate in the PLINK estimates compared to BCFtools Likelihoods. Again using our simulated data as a model, an individual with a true *F*_ROH_ value of 0.40 would be estimated to have an *F*_ROH_ of 0.46 (difference = 0.06) when estimated at 10X-50X with PLINK.

For the two BCFtools methods, patterns of underestimation were consistent with these approaches’ high false negative rates and low false positive rates (Fig. 3). Conversely, PLINK produced higher false positive rates and lower false negative rates than either BCFtools method, consistent with overestimation of *F*_ROH_. In terms of absolute difference between true and called *F*_ROH_ values, PLINK outperformed BCFtools at 10X coverage and above, suggesting that PLINK will often provide the most robust estimate of *F*_ROH_. However, at lower coverages (5X-10X), BCFtools Genotypes could be considered, given that this method produces *F*_ROH_ estimates closer to true *F*_ROH_ than either PLINK or BCFtools Likelihoods. On the other hand, the underestimates produced by this approach are likely related to the high false negative rates we observed (especially relative to PLINK), and the appearance of convergence on true *F*_ROH_ may be due to length-specific ROH calling rates by this program (see below) and therefore highly variable across populations. It is important to note that while the trends we describe may be consistent with some empirical results (*e.g.*, [5]), individual variation in genomic characteristics exerts strong influence over *F*_ROH_ inference results as suggested by comparisons between our simulated and empirical results.

### Demographic history influences F_ROH_ value accuracy

The relationship between true and inferred *F*_ROH_ values varied with demographic scenario, demonstrating the potential for uncertainty when incorporating these inbreeding values into management action plans. For example, in the large population where true *F*_ROH_ values are small (< 0.3), inferred *F*_ROH_ accuracy decreased with increasing values of true *F*_ROH_ for the BCFtools methods, with both tending to underestimate *F*_ROH_ with increasing true *F*_ROH_ (Fig. 2A,E). The opposite trend occurred for the declining population, where true *F*_ROH_ values exceed 0.9, with all three methods approaching more accurate *F*_ROH_ estimates with increasing values of true *F*_ROH_ (Fig. 2D,H,L). These contrasting patterns are likely due to different ROH length distributions in these two demographic scenarios. In the declining population, ROHs in the individuals with the highest overall *F*_ROH_ values tend to comprise very long ROHs (Fig. 1I). Consistent overestimation of *F*_ROH_ for these very long ROHs by BCFtools Genotypes and PLINK directly leads to higher inferred *F*_ROH_ estimates in these individuals. However, individuals with lower overall *F*_ROH_ values have a greater proportion of their ROHs in shorter length categories, which are underestimated by all three methods (Fig. 4D), leading to inferred overall *F*_ROH_ estimates that diverge from true *F*_ROH_ as true *F*_ROH_ decreases (Fig. 2D,H,L). In the large population, although some individuals have higher true *F*_ROH_ values, most ROHs are still short (Fig. 1C), and likely to be underestimated by all three methods. Therefore, as true *F*_ROH_ increases, so do the number of true short ROHs and the cumulative effects of underestimation across methods, leading to overall *F*_ROH_ estimates that diverge from true *F*_ROH_ as true *F*_ROH_ increases.

Overall, we found that the accuracy of ROH detection significantly varies across populations with different demographic histories. Our analysis revealed that declining populations exhibited the highest variance in *F*_ROH_ estimation relative to true *F*_ROH_ values. In contrast, large populations tended to show more stable estimates but with a general tendency for underestimation of true *F*_ROH_, particularly as true *F*_ROH_ values increased. These findings highlight the need for an analysis approach that considers demographic history of the population. For conservationists and researchers working with genomic data from populations known or suspected to be in decline, it is vital to interpret ROH analysis outputs with the breadth of demographic scenarios in mind. When demographic history is unknown and cannot be estimated, the distribution of *F*_ROH_ present in the dataset could be compared to the distributions simulated here (Fig. 1B,D,F,H), or to simulated distributions of *F*_ROH_ with more similar natural history parameters. Although tedious, these approaches will minimize the effects of misestimation of *F*_ROH_, reducing the risk of exacerbating genetic diversity loses in populations and species that are most at risk to extinction.

### Coverage ≤ 10X strongly influences called ROH lengths

For the empirical data at 5X coverage, relative to higher coverage levels, all methods consistently produced lower *F*_ROH_ estimates for short ROHs and higher *F*_ROH_ estimates for longer ROHs (Fig. 7). The overcalling of intermediate to very long ROHs at 5X could be related to the ROH-lumping issue noted in the simulated results, wherein multiple true ROHs are erroneously called as a single ROH (Fig. 6). While we cannot confirm the accuracy of *F*_ROH_ inference for the empirical data, comparisons between results generated at 5X and higher levels of coverage are consistent with the simulated results, suggesting that these patterns are accurate (Fig. 2). For the Tasmanian devil samples we analyzed, the results from 5X coverage suggest much more frequent, recent inbreeding than the results from ≥ 10X coverage, painting a much more dire demographic scenario than is presented when more coverage is obtained. Therefore, if one of the goals of a whole-genome sequencing project is to assess recent or historical patterns of inbreeding from ROH lengths, ∼10X coverage appears to be a minimum requirement for generating robust ROH inferences.

### Patterns of under- or overestimation may vary with ROH length distributions

In our simulated and empirical data, we observed patterns indicating that underlying ROH length distributions influence the patterns of *F*_ROH_ under- and overestimation. For example, even though PLINK produced higher *F*_ROH_ estimates than both BCFtools methods for the simulated data and PLINK and BCFtools Genotypes produced statistically indistinguishable estimates for the empirical data (Fig. 2), length-specific *F*_ROH_ estimates suggest that differences in underlying true ROH length distributions between the simulated and empirical data may be responsible for the differences in relative *F*_ROH_ results we observed. For the simulated data, BCFtools Genotypes increasingly overestimated *F*_ROH_ as ROH length increased (Fig. 5, Fig. S2), with increasing numbers of true ROHs erroneously combined into single called ROHs (Fig. 6). Although we cannot know the true ROH length distributions for the empirical data, long ROHs were called at higher frequencies in the empirical data relative to the simulated data. The tendency of BCFtools Genotypes to overestimate *F*_ROH_ for long ROHs combined with the presence of more called long ROHs in our empirical data set may have minimized differences in overall *F*_ROH_ estimates between BCFtools Genotypes and PLINK in the empirical results relative to the simulated results (Fig. 2). Increased frequencies of long ROHs in the empirical data may have also led to greater differences in *F*_ROH_ between 5X and 10X across all three methods for the empirical results compared to the simulated results (Fig. 2). All three methods call significantly more intermediate to very long ROHs from the empirical data at 5X than at 10X (Fig. 7B-D), and this may be related to the increased false positive rates we noted at 5X in the simulated data. These results again illustrate the effects of a population’s or individual’s actual ROH complement, which is determined by typically unknown demographic and breeding patterns, on the relative reliability and utility of ROH identification programs.

Particularly for endangered species with potentially complicated demographic histories, interpreting ROH patterns in a population may be most accurate when multiple tools are used to create an integrated picture. For example, comparing overall and length-specific *F*_ROH_ estimates between BCFTools/RoH and PLINK can be used to understand the underlying length distributions; an abundance of shorter ROHs would be indicated by higher overall *F*_ROH_ estimates in PLINK compared to BCFtools Genotypes but similar length-specific *F*_ROH_ patterns, whereas a ROH complement comprising many longer ROHs would be indicated by similar overall *F*_ROH_ estimates between PLINK and BCFtools Genotypes but higher intermediate to very long *F*_ROH_ estimates from BCFtools Genotypes related to PLINK. These accurate assessments of past and ongoing inbreeding could then be used to inform management options, such as translocations to ameliorate close inbreeding.

### Conclusions

Inferring the presence and characteristics of ROHs can shed important light on population demographic histories, detect inbreeding depression when combined with fitness information, and even disentangle the mechanisms underlying or loci contributing to inbreeding depression. However, given the variation in ROH-calling accuracy demonstrated here, we caution against direct comparisons of *F*_ROH_ values generated from different data types or sources or using different inference parameters. Data from disparate studies could be combined and re-analyzed in a standardized fashion, although special attention should be paid to variation in reference genome assembly quality for interspecific comparisons [49]. Regardless of the number of data sets to be analyzed, we strongly recommend that studies relying on ROH inference (i) employ at least two ROH-calling programs and interpret their results with each method’s biases in mind and (ii) compare multiple parameter value combinations via sensitivity analysis, taking care to vary parameters of particular relevance to a data set.

## Acknowledgments

We thank members of the Willoughby lab for constructive comments on earlier versions of this manuscript as well as M Kardos and 2 anonymous reviewers.

## Data Availability

Code for all bioinformatic analyses available at https://github.com/avril-m-harder/roh_inference_testing and https://github.com/kennethb22/roh_parameter_project_kk. All FASTA files for simulated individuals and final VCF files (for simulated and empirical data and all coverage levels) will be uploaded to a public repository upon manuscript acceptance.

## Author Contributions

AMH and JRW conceived the study. AMH, KBK, and SM performed formal data analyses. AMH wrote the manuscript with input and final approval from KBK, SM, and JRW.

