## Supplementary material for "Detectability of runs of homozygosity is influenced by analysis parameters and population-specific demographic history": Fig. S

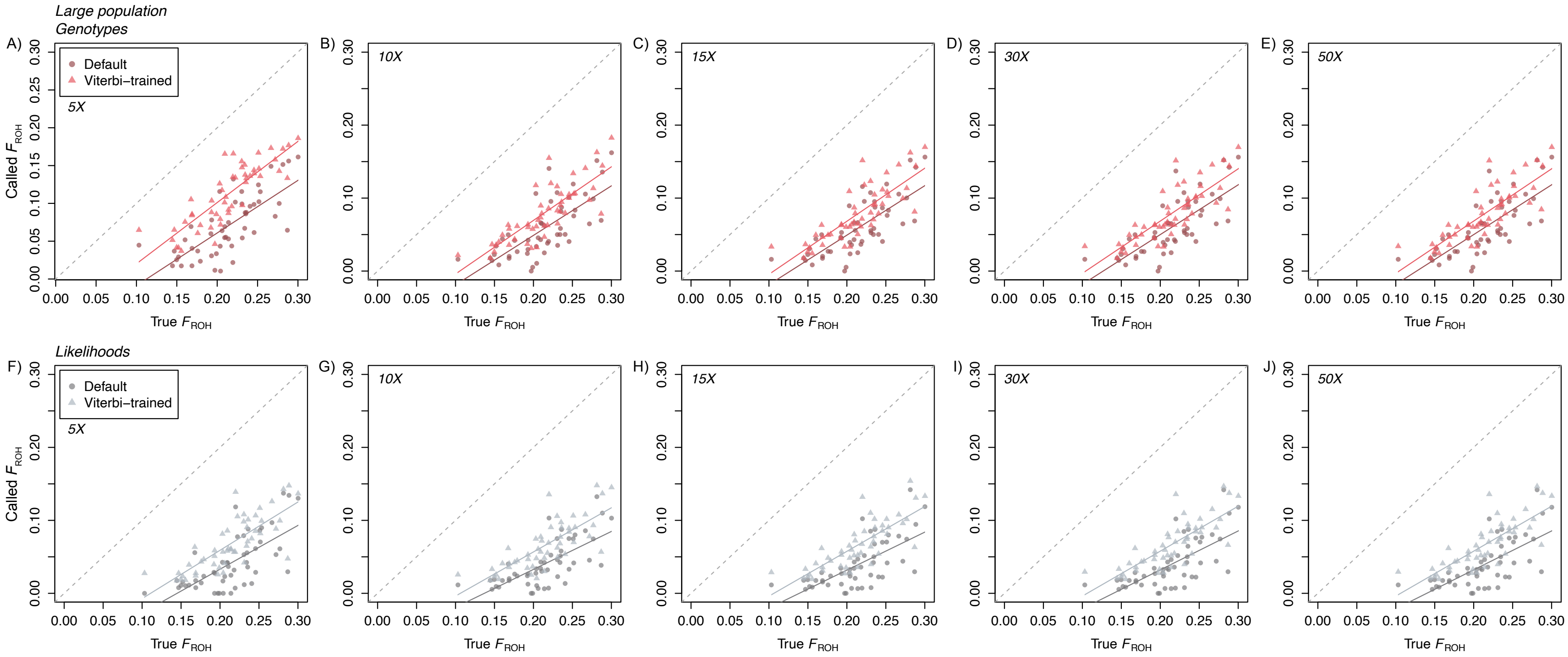


**Figure S1.** True $F_{ROH}$ vs. called $F_{ROH}$ across coverage levels and BCFtools methods for the large population demographic scenario using default HMM transition probability values and Viterbi-trained values. Solid lines indicate linear regression results and the dashed line is the 1:1 line. Within each method and coverage level combination, the slope and intercept parameters did not significantly differ between the approaches using Viterbi-trained or default probabilities, but the regression line calculated using Viterbi-trained results was consistently closer to the 1:1 line.

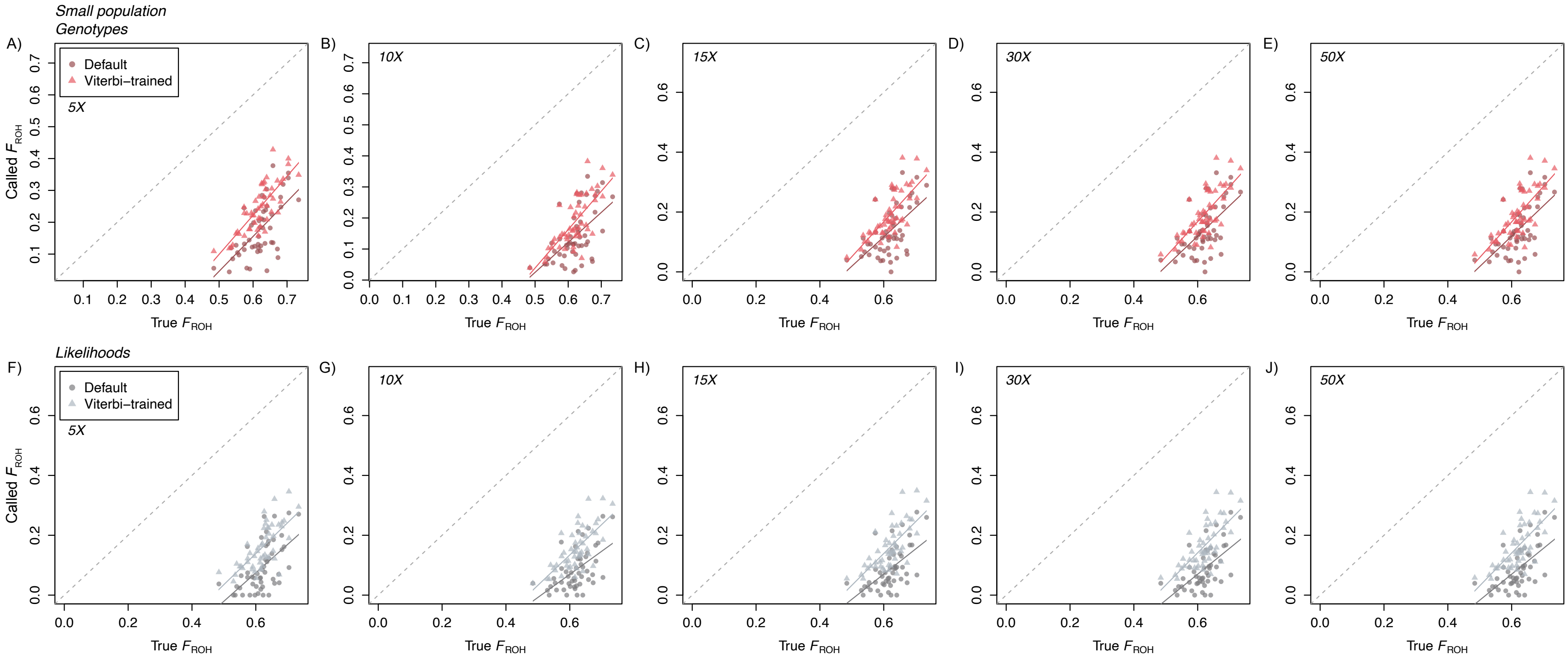


**Figure S2.** True $F_{ROH}$ vs. called $F_{ROH}$ across coverage levels and BCFtools methods for the small population demographic scenario using default HMM transition probability values and Viterbi-trained values. Solid lines indicate linear regression results and the dashed line is the 1:1 line. Within each method and coverage level combination, the slope and intercept parameters did not significantly differ between the approaches using Viterbi-trained or default probabilities, but the regression line calculated using Viterbi-trained results was consistently closer to the 1:1 line.

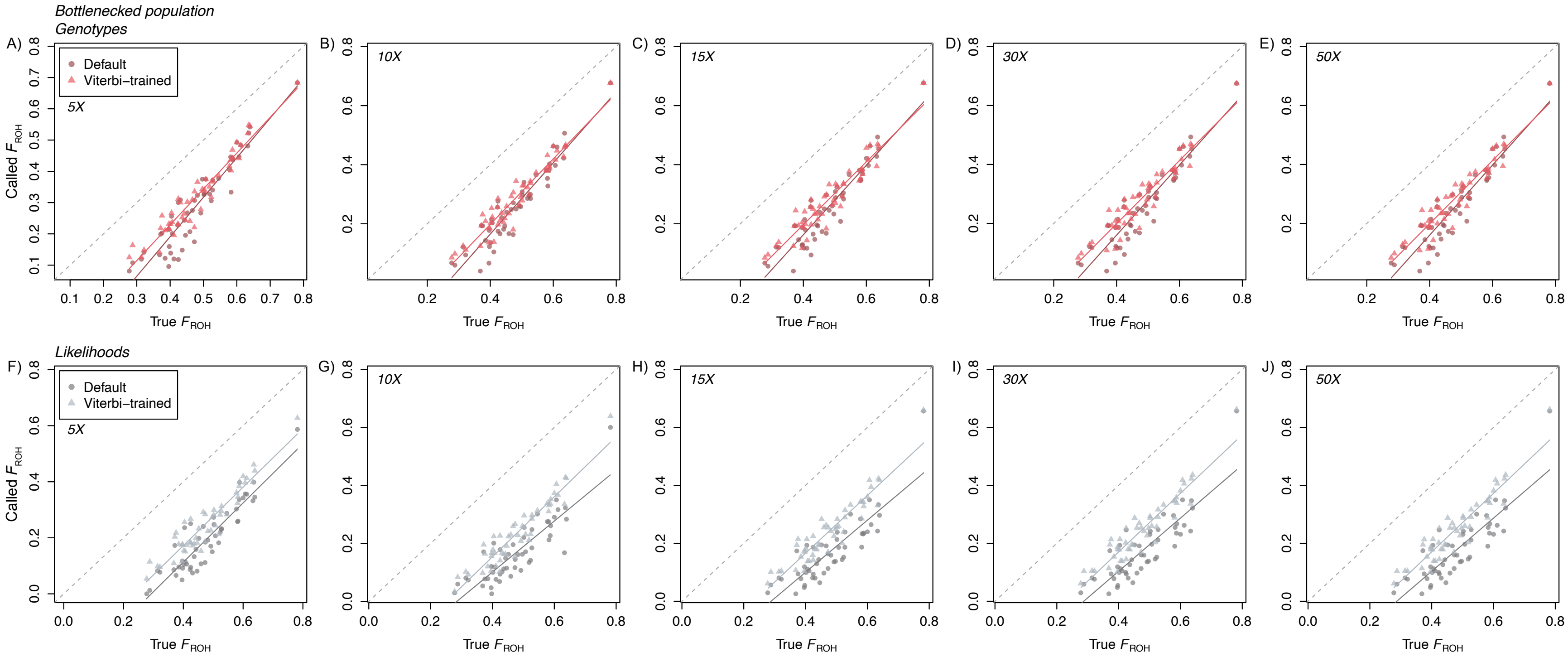


**Figure S3.** True $F_{\mathrm{ROH}}$ vs. called $F_{\mathrm{ROH}}$ across coverage levels and BCFtools methods for the bottlenecked population demographic scenario using default HMM transition probability values and Viterbi-trained values. Solid lines indicate linear regression results and the dashed line is the 1:1 line. Within each method and coverage level combination, the slope and intercept parameters did not significantly differ between the approaches using Viterbi-trained or default probabilities, but the regression line calculated using Viterbi-trained results was consistently closer to the 1:1 line.

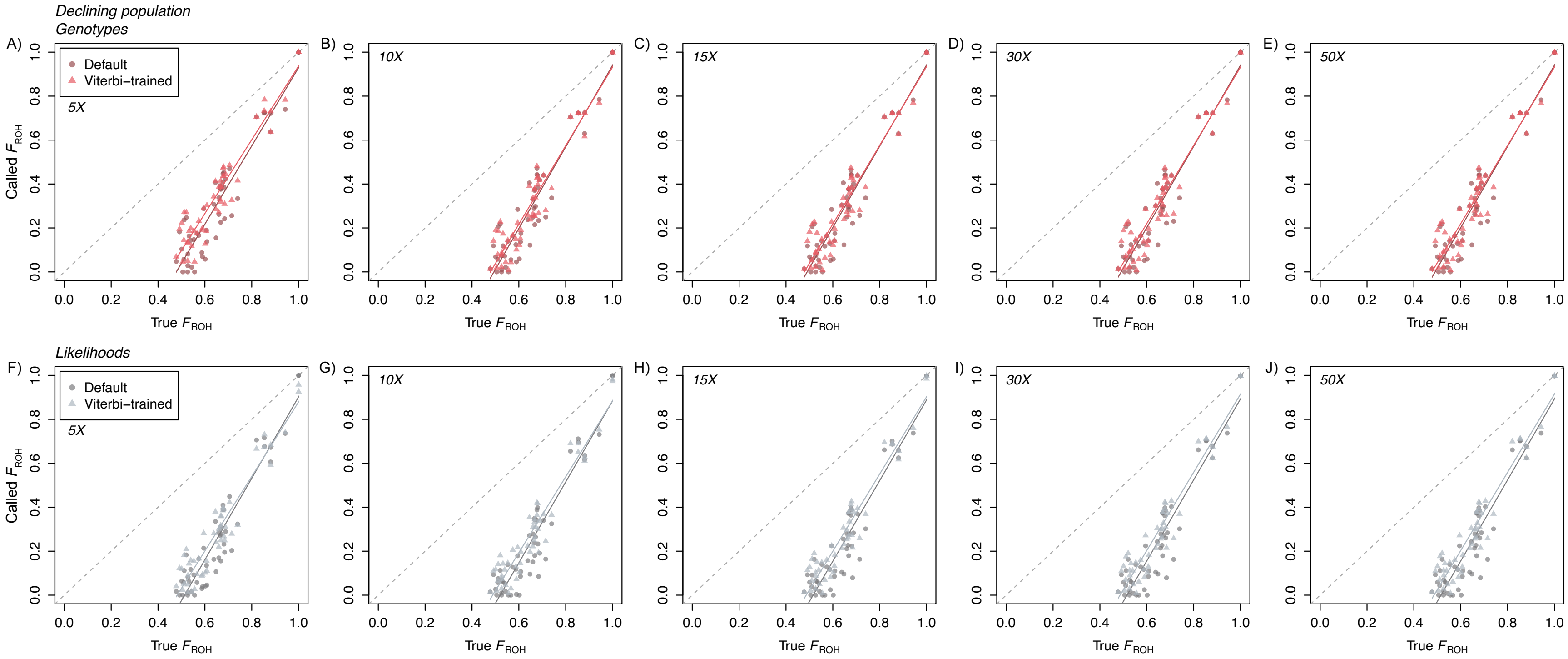


**Figure S4.** True $F_{ROH}$ vs. called $F_{ROH}$ across coverage levels and BCFtools methods for the declining population demographic scenario using default HMM transition probability values and Viterbi-trained values. Solid lines indicate linear regression results and the dashed line is the 1:1 line. Within each method and coverage level combination, the slope and intercept parameters did not significantly differ between the approaches using Viterbi-trained or default probabilities, but the regression line calculated using Viterbi-trained results was consistently closer to the 1:1 line.

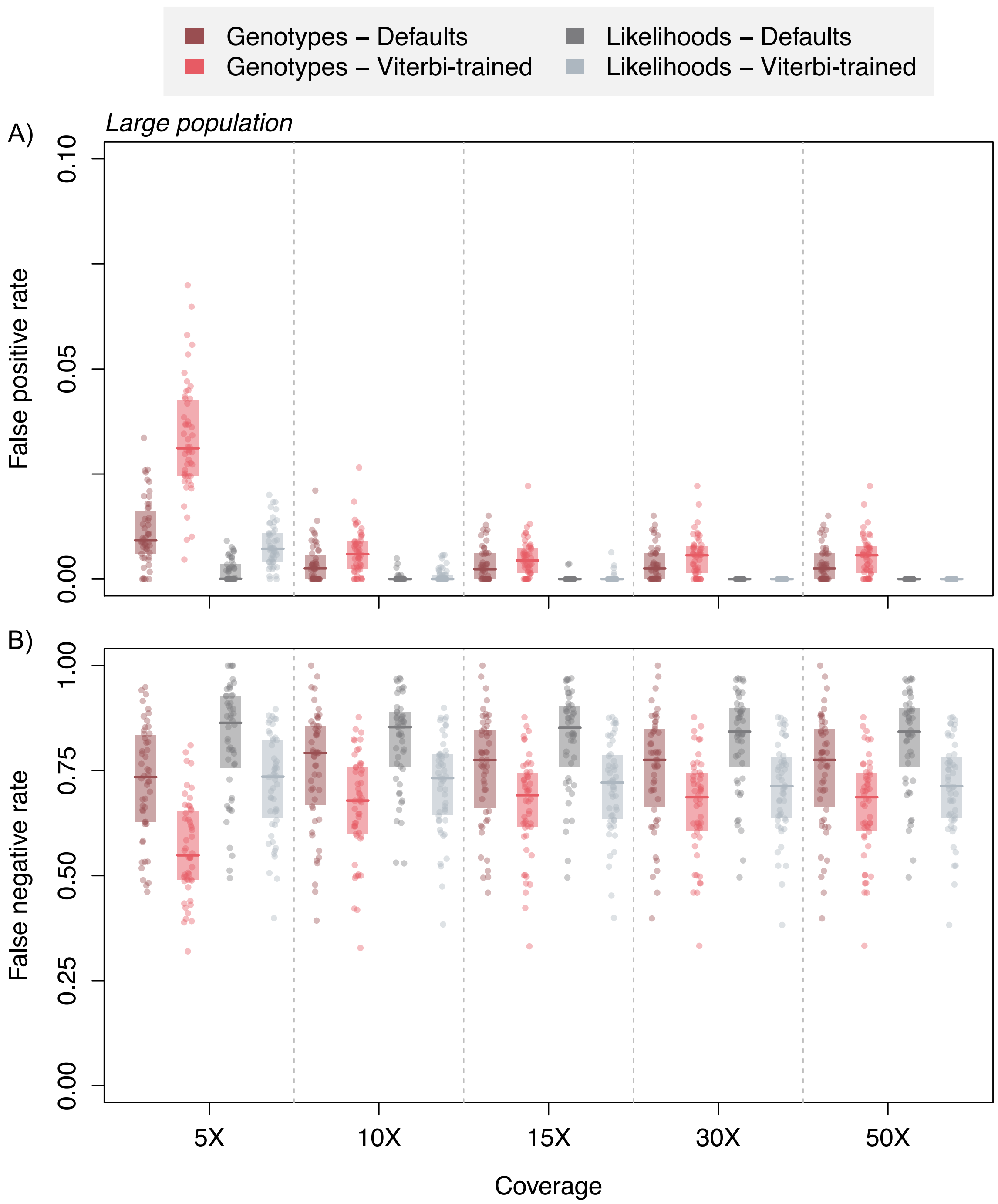


**Figure S5.** A) False positive (*i.e.*, incorrectly calling a base position as being located in a ROH) and B) false negative (*i.e.*, failing to identify a base position as being located in a ROH) rates across coverage levels for the large population demographic scenario. Horizontal lines indicate median values and shaded boxes are 50% quantiles. Note difference in scale of *y*-axis between panels A and B. Although not significantly different, rates calculated for ROHs called using Viterbi-trained HMM transition probabilities tend to be lower than rates calculated for ROHs called using default probabilities.

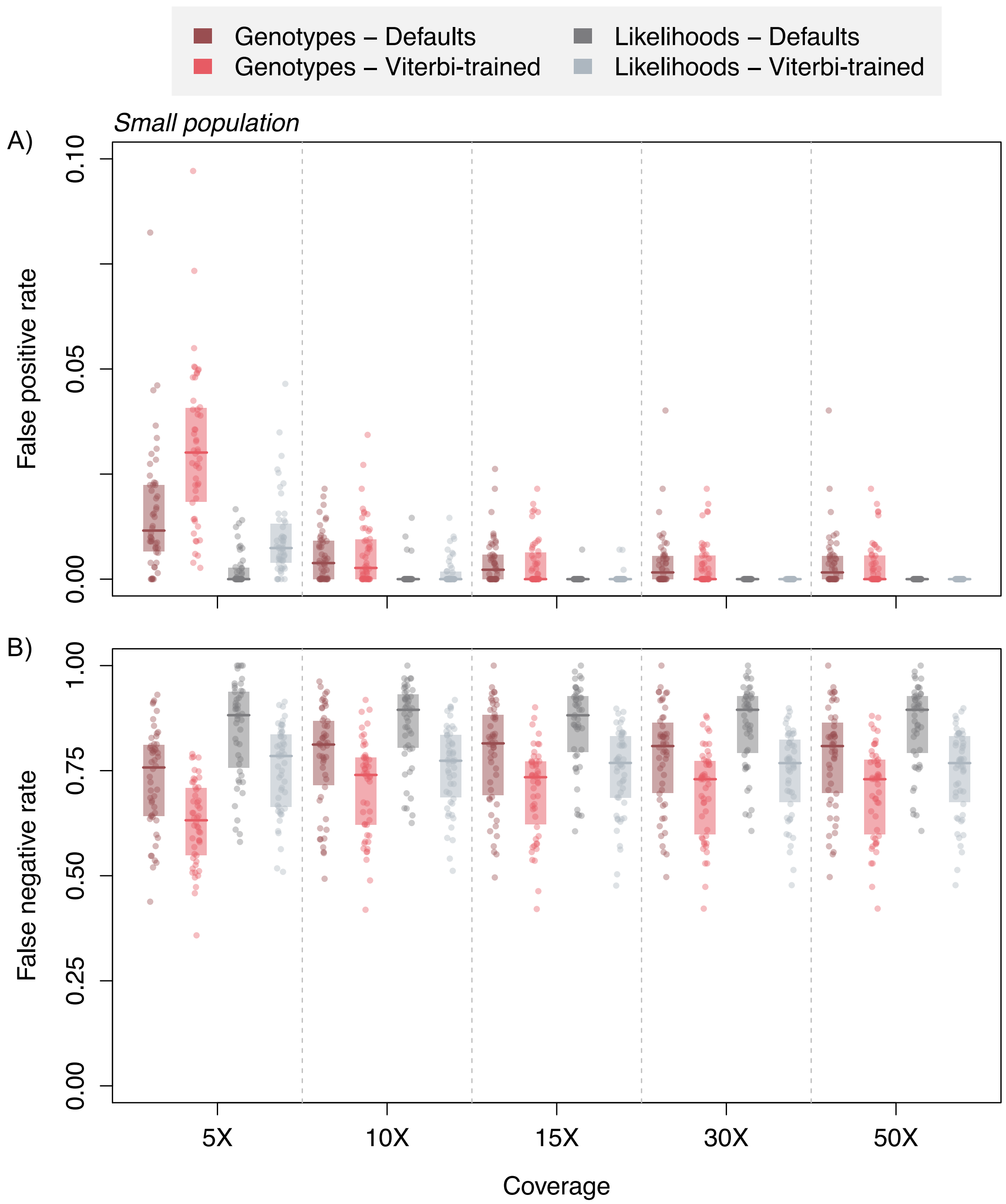


**Figure S6.** A) False positive (*i.e.*, incorrectly calling a base position as being located in a ROH) and B) false negative (*i.e.*, failing to identify a base position as being located in a ROH) rates across coverage levels for the small population demographic scenario. Horizontal lines indicate median values and shaded boxes are 50% quantiles. Note difference in scale of *y*-axis between panels A and B. Although not significantly different, rates calculated for ROHs called using Viterbi-trained HMM transition probabilities tend to be lower than rates calculated for ROHs called using default probabilities.

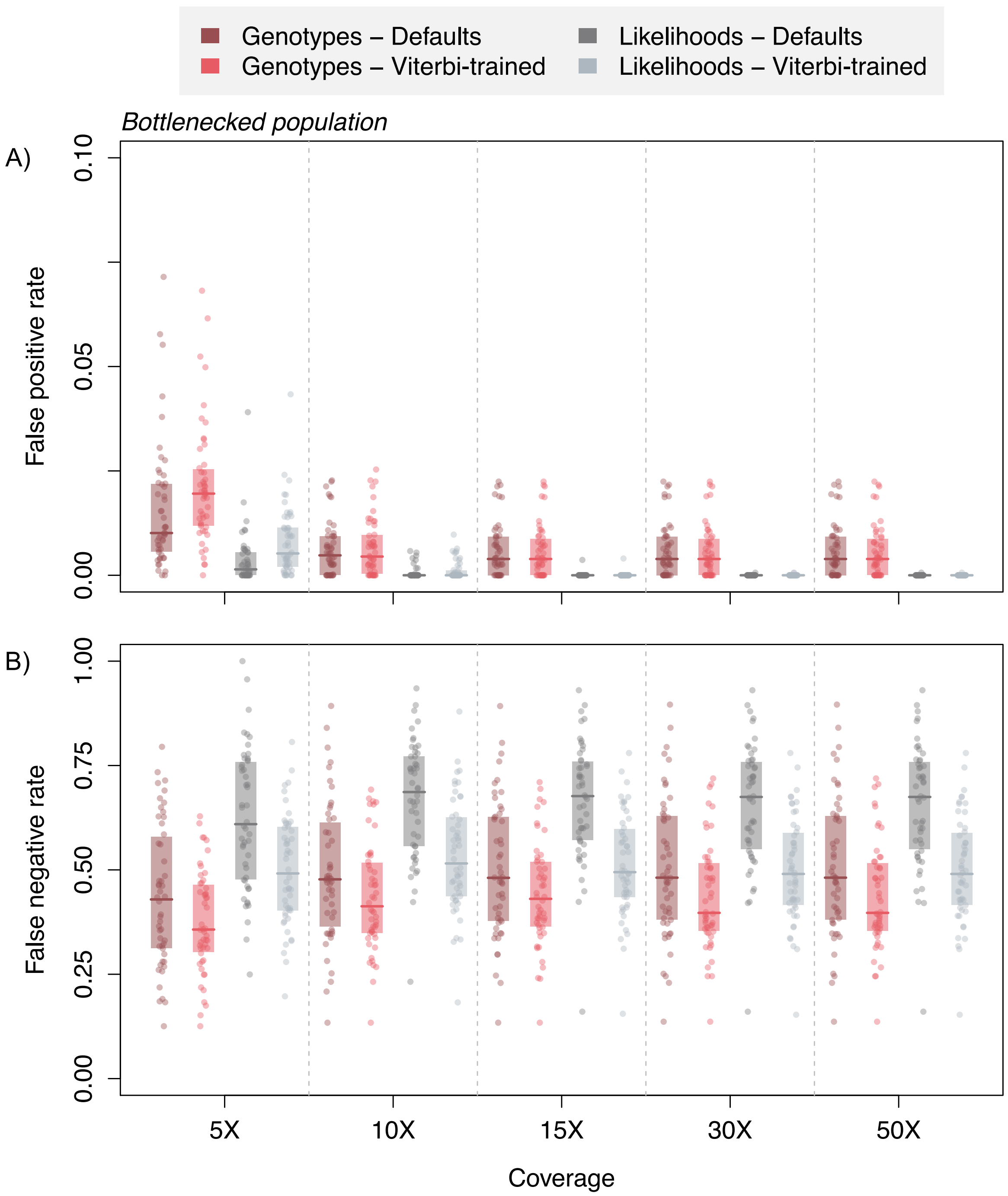


**Figure S7.** A) False positive (*i.e.*, incorrectly calling a base position as being located in a ROH) and B) false negative (*i.e.*, failing to identify a base position as being located in a ROH) rates across coverage levels for the bottlenecked population demographic scenario. Horizontal lines indicate median values and shaded boxes are 50% quantiles. Note difference in scale of *y*-axis between panels A and B. Although not significantly different, rates calculated for ROHs called using Viterbi-trained HMM transition probabilities tend to be lower than rates calculated for ROHs called using default probabilities.

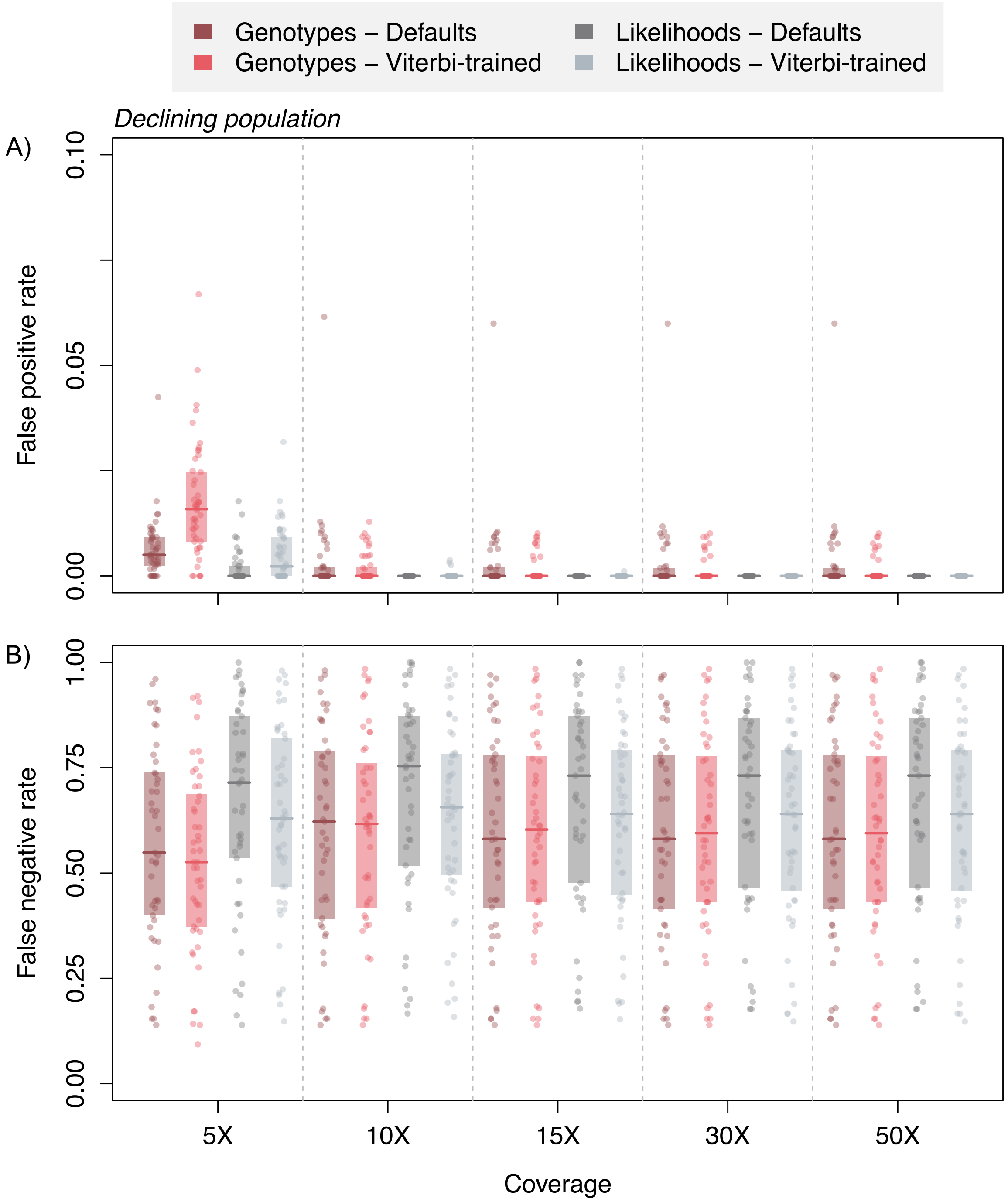


**Figure S8.** A) False positive (*i.e.*, incorrectly calling a base position as being located in a ROH) and B) false negative (*i.e.*, failing to identify a base position as being located in a ROH) rates across coverage levels for the declining population demographic scenario. Horizontal lines indicate median values and shaded boxes are 50% quantiles. Note difference in scale of *y*-axis between panels A and B. Although not significantly different, rates calculated for ROHs called using Viterbi-trained HMM transition probabilities tend to be lower than rates calculated for ROHs called using default probabilities.

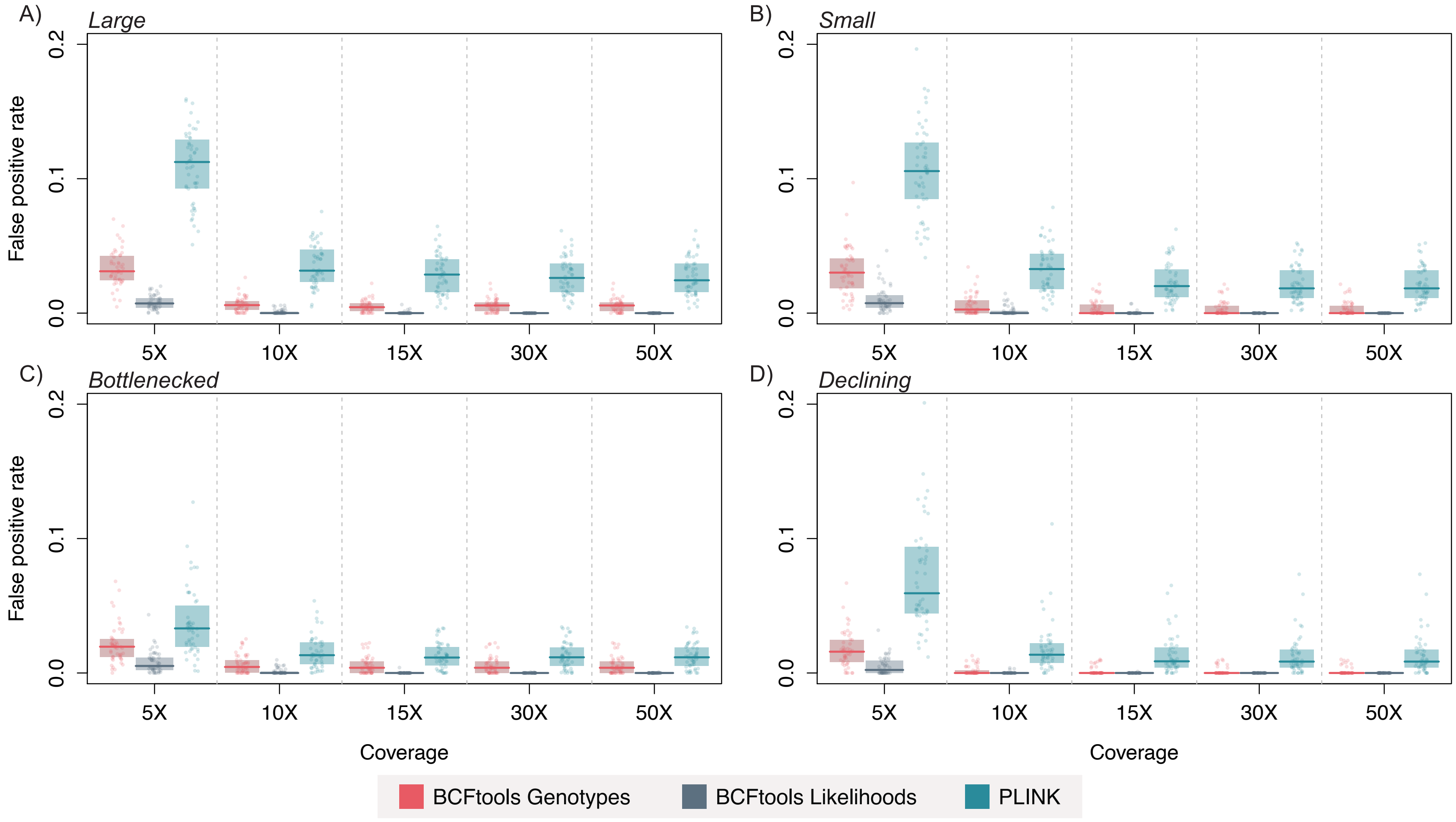


**Figure S9.** False positive (*i.e.*, incorrectly calling a base position as being located in a ROH) rates across coverage levels for all population demographic scenarios. Horizontal lines indicate median values and shaded boxes are 50% quantiles.

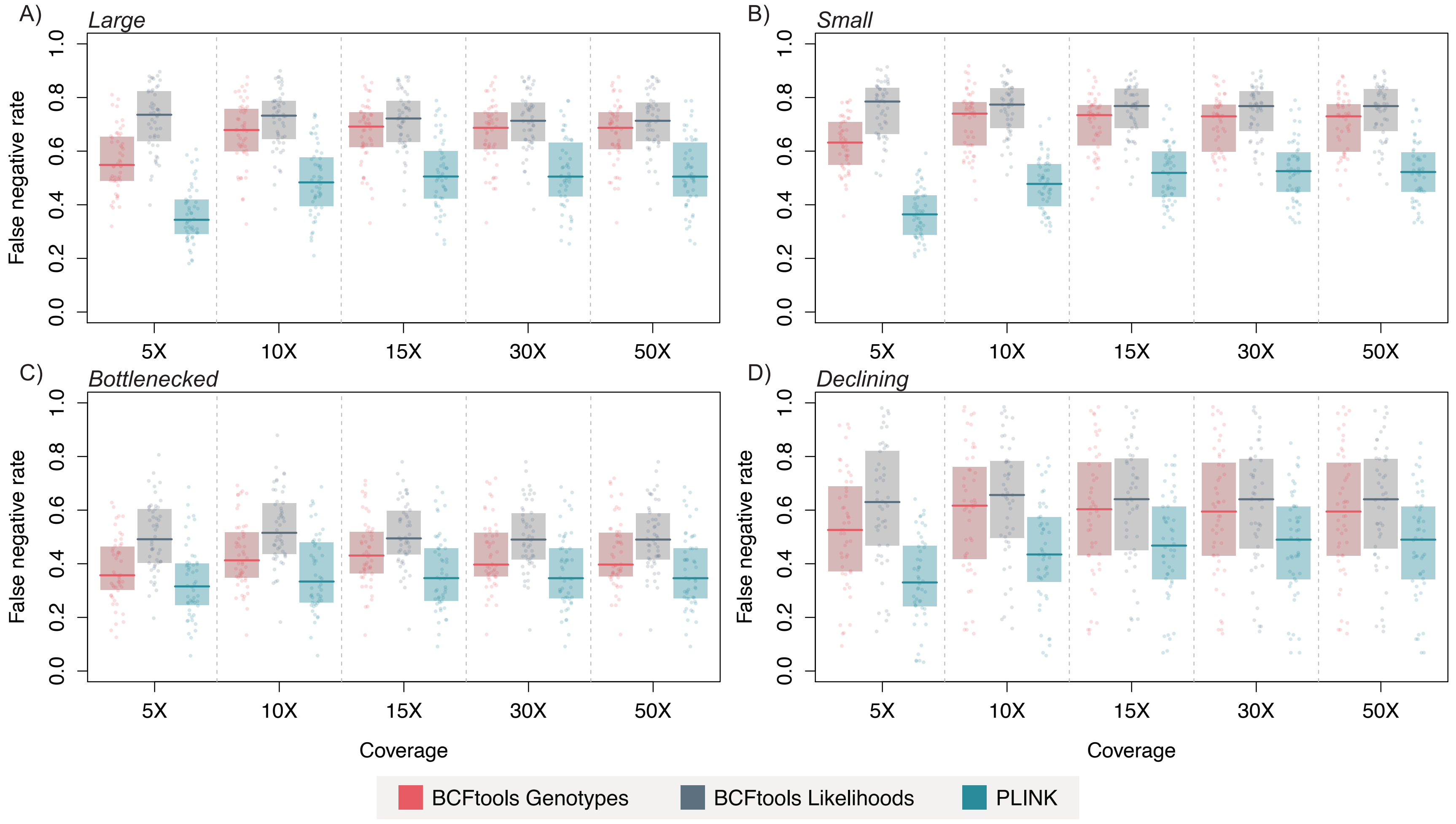


**Figure S10.** False negative (*i.e.*, failing to identify a base position as being located in a ROH) rates across coverage levels for all demographic scenarios. Horizontal lines indicate median values and shaded boxes are 50% quantiles.

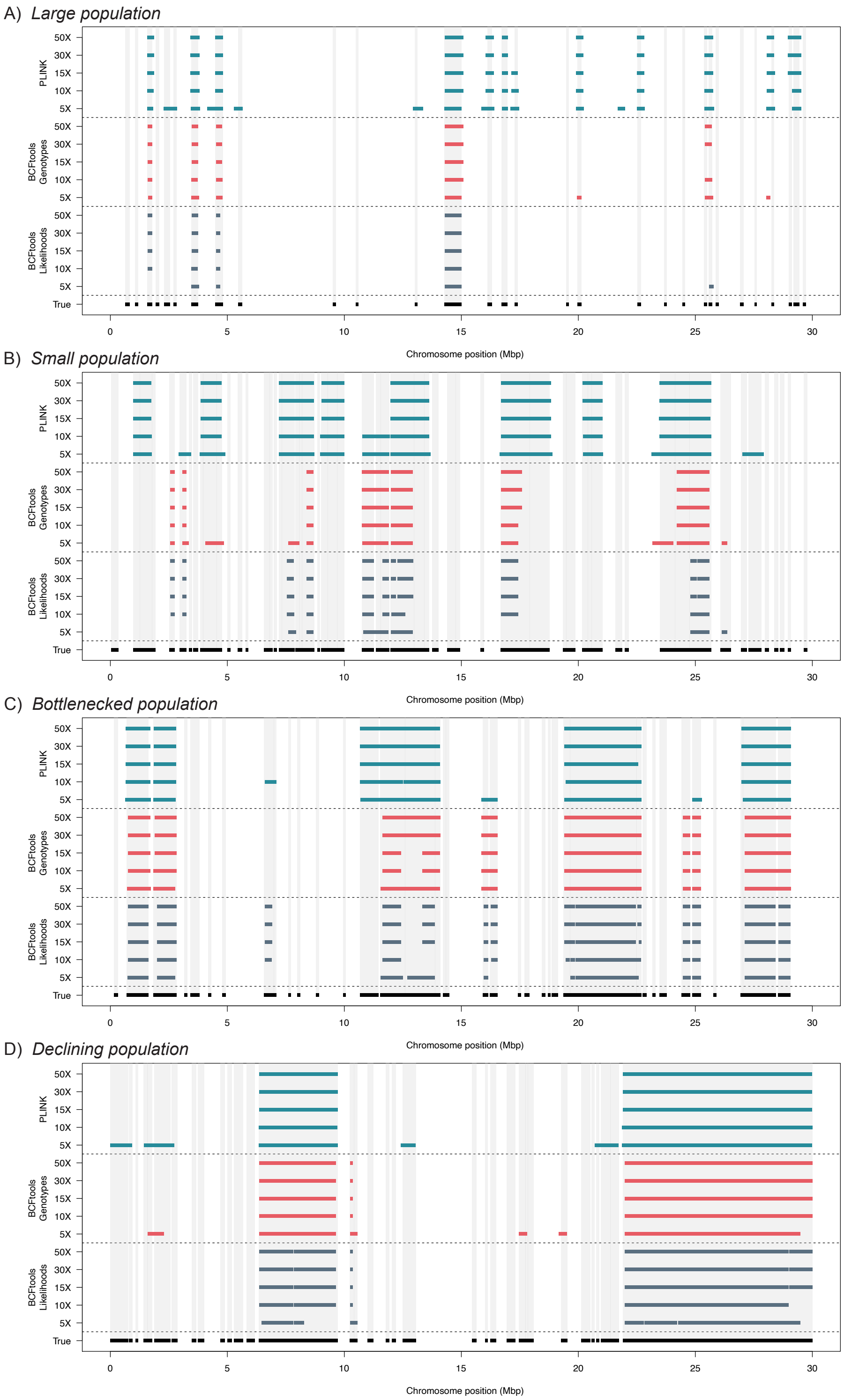


**Figure S11.** True and called ROHs ≥ 100 kb in length for one individual from each demographic scenario simulated. True ROHs are indicated by black polygons and light gray shading.

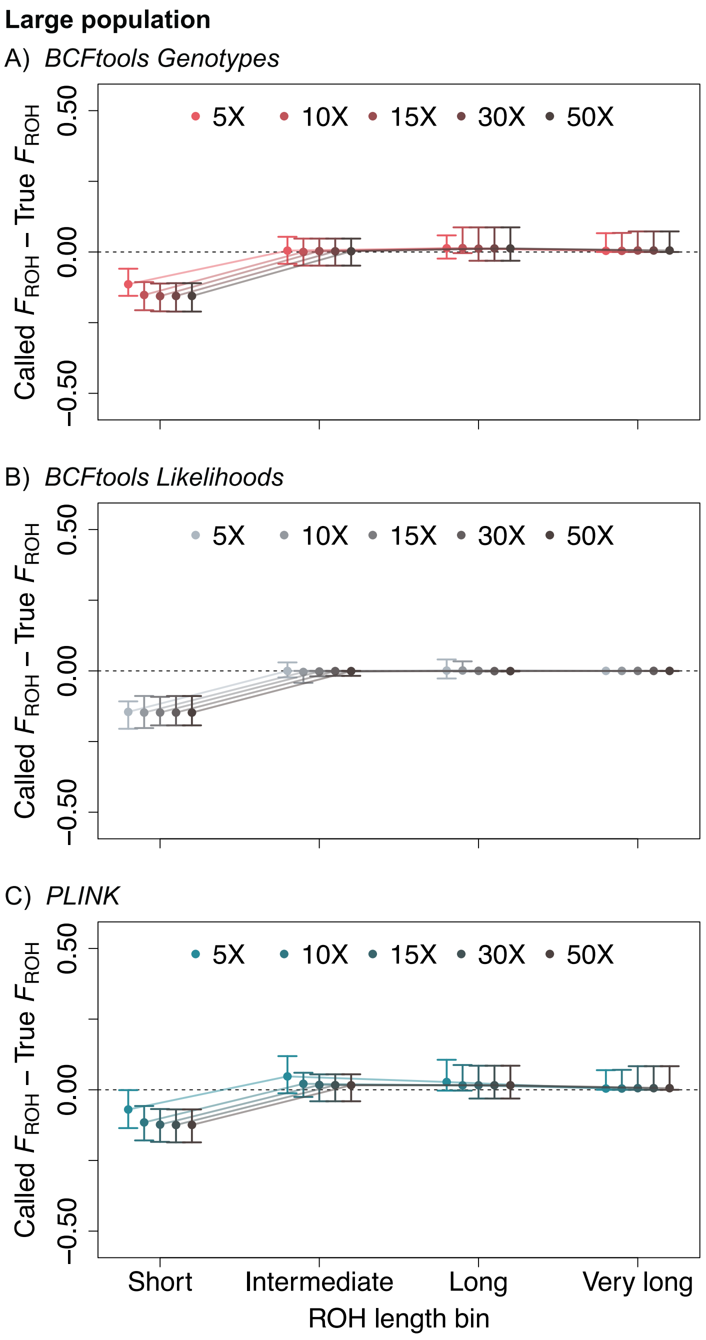


**Figure S12.** Called $F_{\mathrm{ROH}}$ – True $F_{\mathrm{ROH}}$ by length bin and coverage level for the large population demographic scenario.

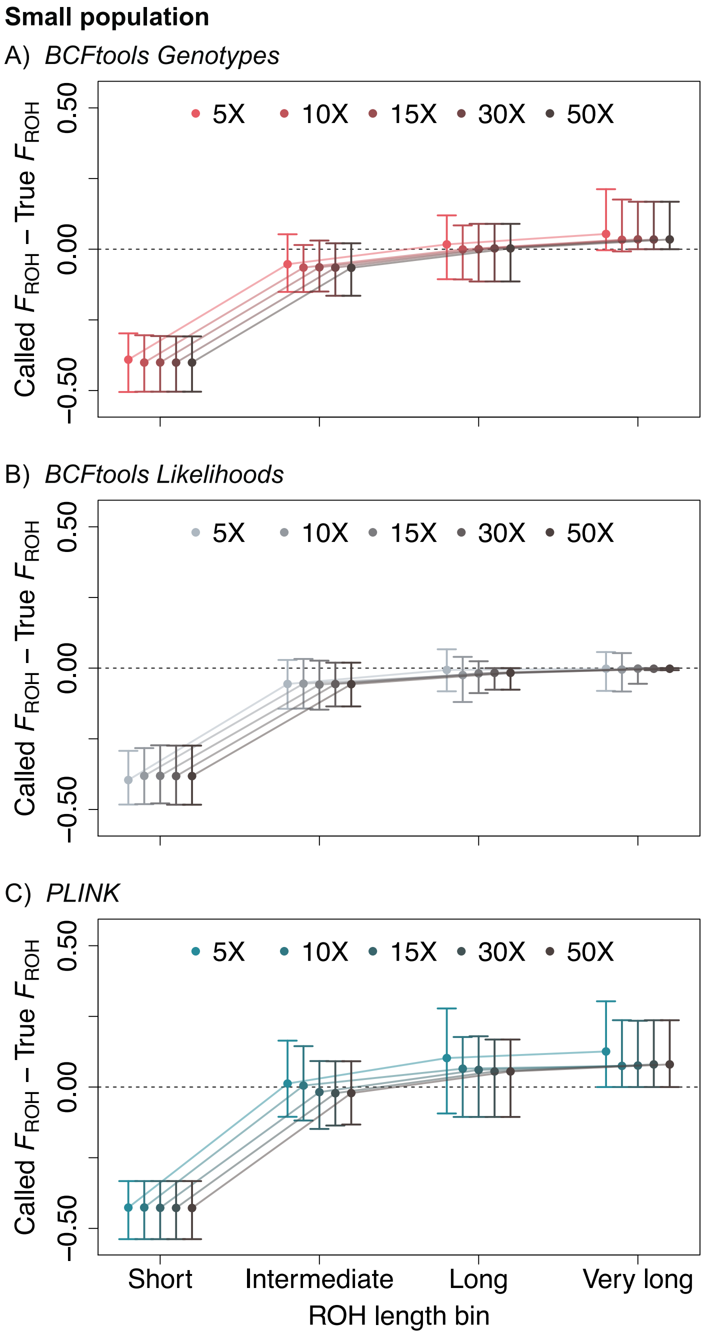


**Figure S13.** Called $F_{\mathrm{ROH}}$ – True $F_{\mathrm{ROH}}$ by length bin and coverage level for the small population demographic scenario.

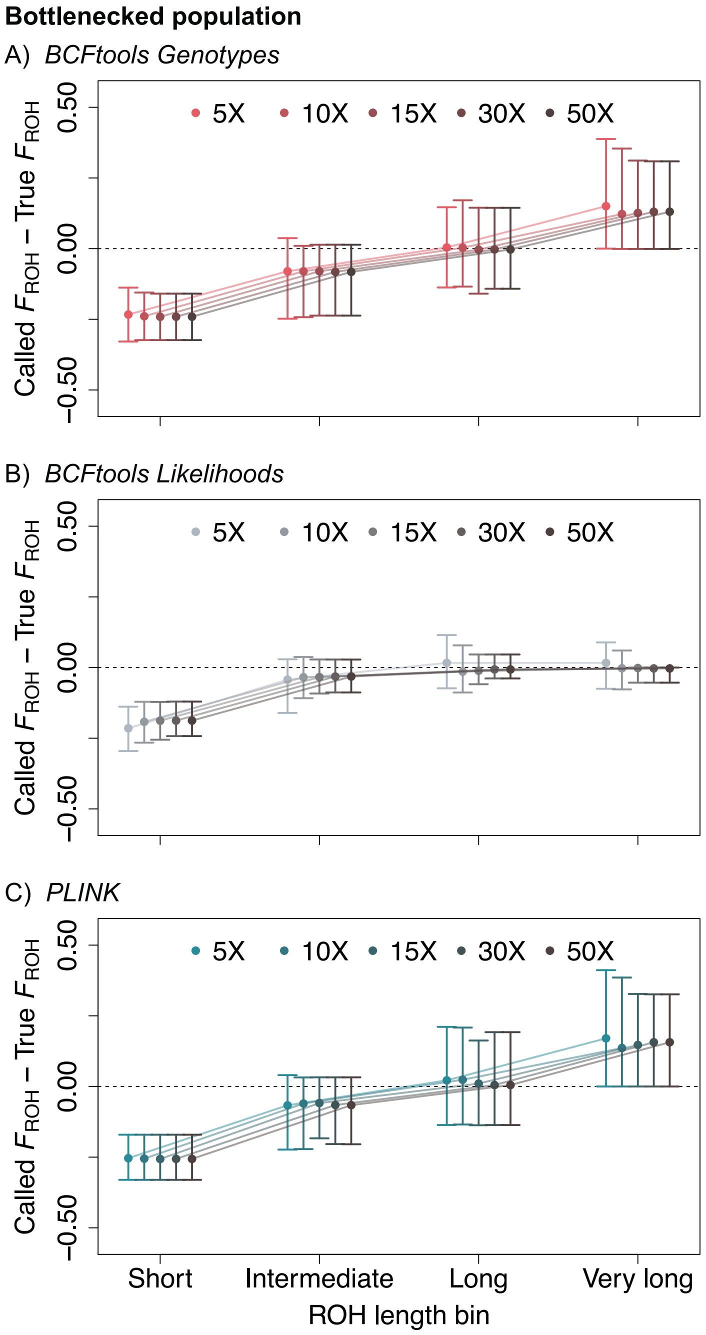


**Figure S14.** Called $F_{ROH}$ – True $F_{ROH}$ by length bin and coverage level for the bottlenecked population demographic scenario.

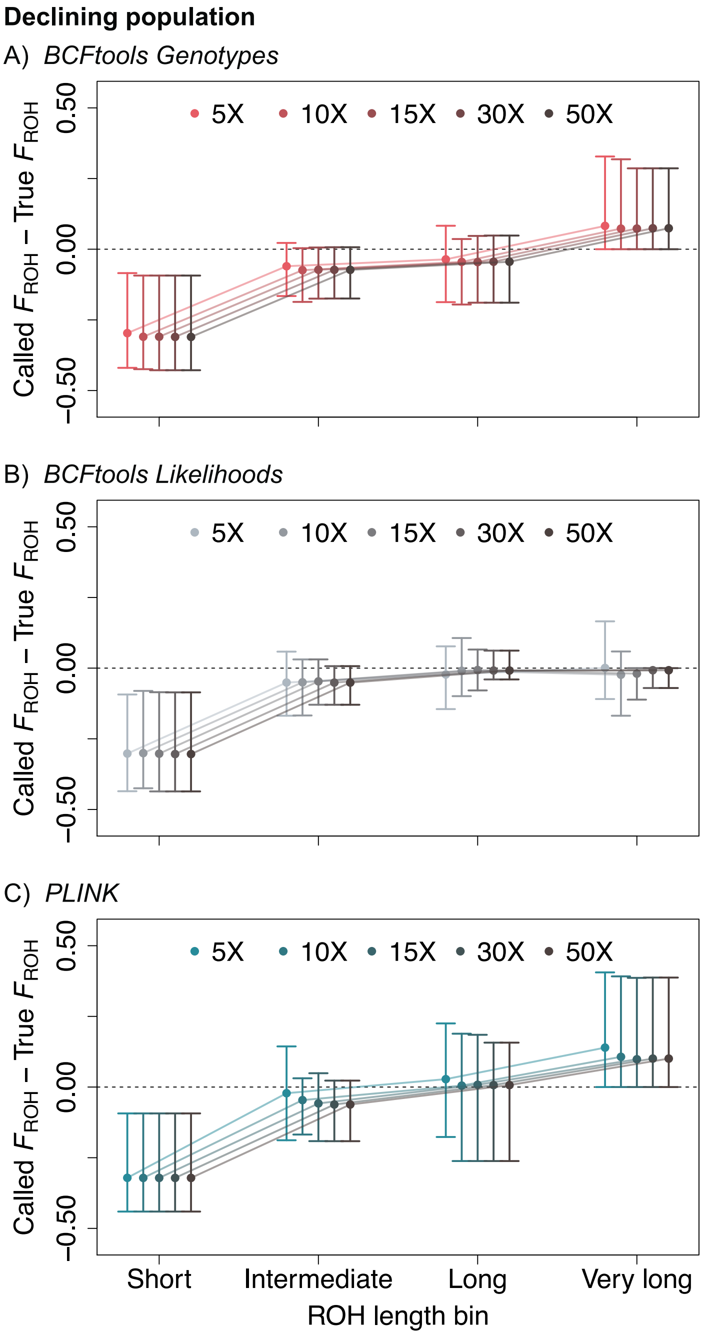


**Figure S15.** Called $F_{\mathrm{ROH}}$ – True $F_{\mathrm{ROH}}$ by length bin and coverage level for the declining population demographic scenario.

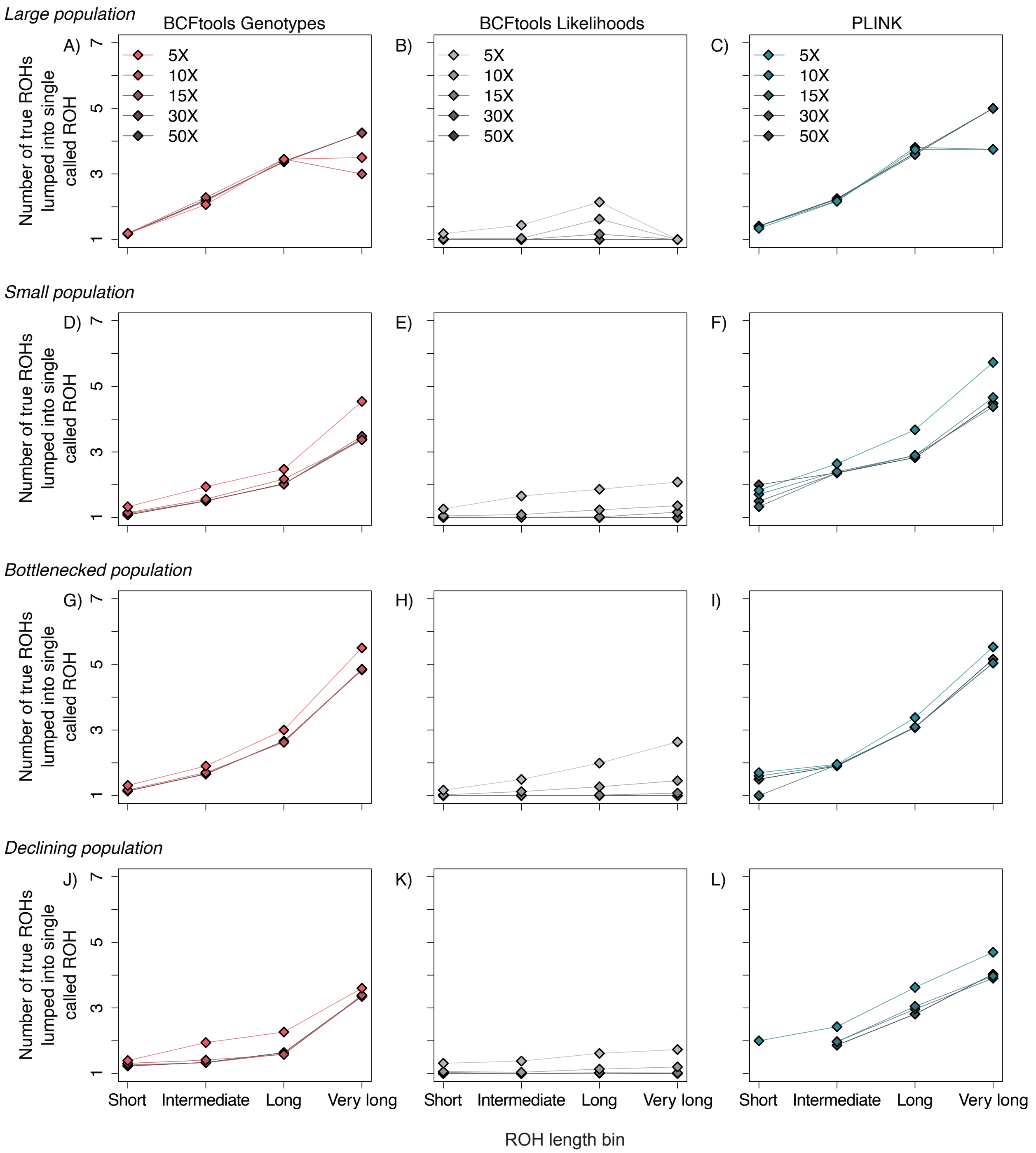


**Figure S16.** Mean number of true ROHs combined into a single called ROH at each coverage level by length bins. Results are presented for all four demographic scenarios.
